# Blocking MIF secretion enhances CAR T-cell efficacy against neuroblastoma

**DOI:** 10.1101/2024.04.05.588098

**Authors:** Josephine G. M. Strijker, Guillem Pascual-Pasto, Yannine J. Kalmeijer, Elisavet Kalaitsidou, Chunlong Zhao, Brendan McIntyre, Stephanie Matlaga, Lindy L. Visser, Marta Barisa, Courtney Himsworth, Rivani Shah, Henrike Muller, Linda G. Schild, Peter G. Hains, Qing Zhong, Roger Reddel, Phillip J. Robinson, Xavier Catena, María S. Soengas, Thanasis Margaritis, Frank J. Dekker, John Anderson, Jan J. Molenaar, Kristopher R. Bosse, Wei Wu, Judith Wienke

**Affiliations:** Princess Máxima Center for Pediatric Oncology, Utrecht, The Netherlands; Division of Oncology and Center for Childhood Cancer Research, The Children’s Hospital of Philadelphia, Philadelphia, Pennsylvania, USA; Singapore Immunology Network (SIgN), Agency for Science, Technology and Research (A*STAR), 8A Biomedical Grove, Immunos, Singapore 138648, Republic of Singapore; Department of Pharmacy and Pharmaceutical Sciences, National University of Singapore, 117543, Singapore, Singapore; Department of Chemical and Pharmaceutical Biology, Groningen, Research Institute of Pharmacy (GRIP), University of Groningen, Antonius Deusinglaan 1, 9713 AV, Groningen, The Netherlands; UCL Great Ormond St Institute of Child Health, London, UK; ProCan, Children’s Medical Research Institute, The University of Sydney, Westmead, NSW, Australia; Melanoma Laboratory, Molecular Oncology Programme, Spanish National Cancer Research Centre (CNIO), Madrid, Spain; Department of Pharmaceutical Sciences, University Utrecht, Utrecht, The Netherlands; Department of Pediatrics; Perelman School of Medicine at the University of Pennsylvania; Philadelphia, PA, 19104; USA; Biomolecular Mass Spectrometry and Proteomics, Bijvoet Center for Biomolecular Research and Utrecht Institute for Pharmaceutical Sciences, Utrecht University, Utrecht, The Netherlands

**Keywords:** Neuroblastoma, CAR T-cell therapy, tumor microenvironment, multi-omics, secretome, MIF, PROTAC

## Abstract

While chimeric antigen receptor (CAR) T-cell therapies are showing highly promising first results in neuroblastoma, immunosuppressive tumor microenvironments (TME) limit T cell persistence and durable clinical efficacy. To improve CAR T-cell efficacy further, we applied a multi-omics approach including single-cell RNA sequencing and proteomics, which identified 13 targetable immunosuppressive factors in neuroblastoma. Of these, macrophage migration inhibitory factor (MIF) and midkine (MDK) were validated across multiple published RNA datasets. Moreover, they were secreted in high abundance by neuroblastoma tumoroids. Functional validation experiments revealed MIF as a potent inhibitor of CAR T-cells, *in vitro* and *in vivo.* Degradation of MIF by PROTAC technology significantly enhanced CAR T-cell activation targeting GPC2 and B7-H3, providing a potential intervention against MIF. By defining the immunosuppressive effects of neuroblastoma’s TME on CAR T-cell efficacy, particularly the pivotal role of MIF, we provide a therapeutic strategy for improving adoptive cell therapies for this pediatric malignancy.

## Introduction

Following major successes in immunotherapies for adult cancer patients, immunotherapies are now also entering the stage in pediatric solid cancers^1–6^. Adoptive cell therapy, which includes chimeric antigen (CAR) T-cell technology, has led to substantially increased survival rates in multiple cancers and has recently also shown early promise in neuroblastoma^3,7,8^.

Neuroblastoma is an extracranial solid tumor arising in the adrenal gland and sympathetic ganglia^9^. Risk factors such as age, metastases and genetic alterations, like *MYCN* amplification, are used to stratify patients into risk groups^10–12^. Particularly the high-risk group has a dismal 5-year survival rate of ∼50%, despite an extensive treatment regimen including surgery, high-dose chemotherapy, radiotherapy, and immunotherapy with Dinutuximab, a monoclonal antibody targeting GD2^13,14^.

CAR T-cell therapy for neuroblastoma had not been convincingly successful, until a recent clinical trial with third-generation GD2 targeting CAR T-cells showed encouraging results^7,15–19^. Nevertheless, clinical responses were limited to patients with lower disease burden, suggesting that larger bulk tumors remain a significant challenge for neuroblastoma CAR T-cell therapies^7^. Several factors may prevent adequate CAR T-cell efficacy in neuroblastoma and other solid tumors, including suboptimal CAR T-cell proliferation and persistence, and limited tumor infiltration^20–22^. Moreover, particularly in solid tumors like neuroblastoma, the immunosuppressive tumor-microenvironment (TME) poses an additional significant challenge for CAR T-cells^6,23,24^. For example, in neuroblastoma, the rate of cytotoxic lymphocyte infiltration into the tumor is lower in patients with high-risk disease and patients with a worse prognosis, clearly suggesting that high-risk neuroblastomas can evade an immune response^25,26^.

The immunosuppressive microenvironment in neuroblastomas include expression of well documented immune checkpoints such as PD-L1 and CD200^24,27–30^. In addition, neuroblastoma cells secrete immunoregulatory mediators such as TGF-β, galectin-1, and soluble GD2^31–35^, that likely hamper proper activation of lymphocytes (and CAR T-cells). Beyond the immediate TME, secreted factors may also reach the systemic circulation, where immunosuppressive effects may be exerted on peripheral lymphocytes. Indeed, peripheral T cells in neuroblastoma patients are reduced in numbers and unresponsive to *in vitro* activation, hinting at a much more widespread dysfunction of T cells in patients with neuroblastoma^36–42^. Such peripheral immunosuppressive effects may also hamper the long-term persistence of CAR T-cells, their migration, as well as their activation capacity.

Currently, several flavors of CAR T-cells are under development for neuroblastoma either in early phase clinical trial or late-phase preclinical testing. Amongst these are a second-generation CAR targeting the signaling co-receptor glypican 2 (GPC2), and a second-generation CAR targeting B7-H3, which is a well-known tumor antigen expressed by multiple pediatric cancers, including neuroblastoma^43,44^.

In order to improve CAR T-cell efficacy, we sought to identify immunosuppressive factors secreted by neuroblastoma that may inhibit CAR T-cells and hamper their function. Using a multi-omics approach combining single-cell RNA sequencing and proteomics, we identified Macrophage Migration Inhibitory Factor (MIF) and Midkine (MDK) as secreted factors that could drive immunosuppression locally and distally in neuroblastoma. Conversely, MIF depletion significantly increased CAR T-cell efficacy *in vitro* and *in vivo*. Taken together, we propose that targeting MIF may be a viable strategy to increase the efficacy of CAR T-cell therapy for patients with neuroblastoma.

## Results

### MIF and MDK are negatively associated with lymphocyte cytotoxicity in neuroblastoma

We recently observed, by single-cell RNA sequencing of 24 neuroblastoma tumors, that natural killer (NK) cells in the neuroblastoma TME have reduced cytotoxic potential, and that T cells in the same TME display features of dysfunction^30^. To precisely identify which specific immunosuppressive factors in the neuroblastoma TME may be contributing to this reduced NK and T cell cytotoxic potential, we performed an interaction analysis using CellChat^45^. We focused on interactions of cytotoxic lymphocytes, i.e. NK, CD8^+^ T cells, and γδ-T cells, with other cells in the TME. To specifically define which of these predicted interactions may reduce the cytotoxic potential of these immune cell subsets, we analyzed which interactions negatively correlated with their expression of genes encoding cytotoxic mediators (Fig. 1a). To this end, we constructed a cytotoxicity score for each immune cell subset using 7 genes (*NKG7*, *CCL5*, *CST7*, *PRF1*, *GZMA*, *GZMB* and *IFNG*; Fig. 1b,c). Correlation of this cytotoxicity score with the predicted interaction partners yielded 23 interactions involving 13 unique genes, all of which negatively correlated with NK, CD8^+^ T and/or γδ-T cell cytotoxicity (Fig. 1d). Remarkably, all of these 13 genes were annotated as predicted secreted factors by the SEPDB database of secreted proteins and could thus potentially affect NK/T cells both locally in the TME and distally^46^.

**Figure 1:**
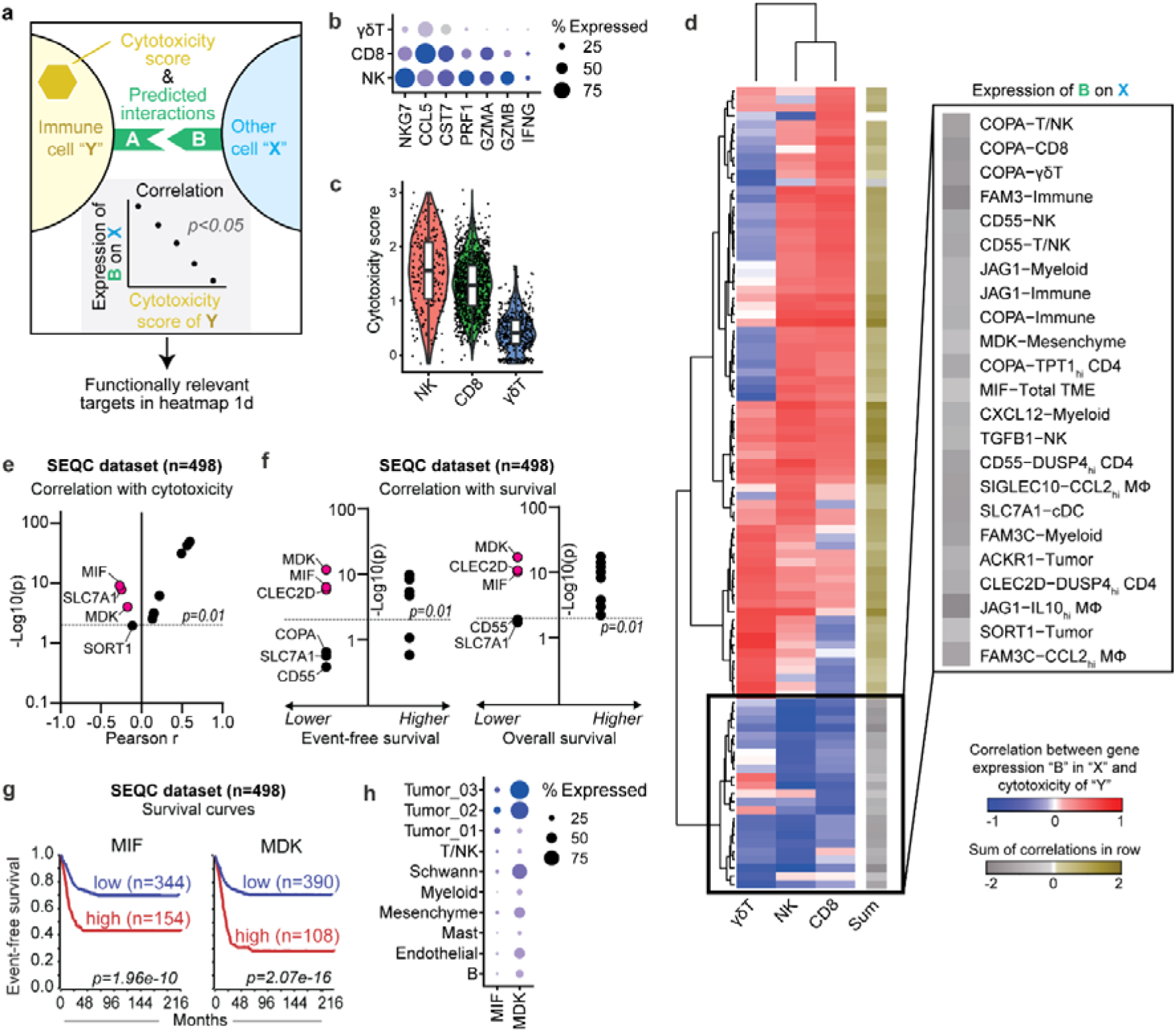
Identification of MIF and MDK as immunosuppressive factors based on transcriptomic data. **a**, Schematic representation of analysis strategy of scRNAseq data, published earlier by our group, in order to determine significant correlations between the cytotoxicity score of immune cell subsets and predicted interactions of those immune cells with other cells in the tumor microenvironment. Adapted from Wienke, et al^30^. **b,** Dotplot representing the gene expression of 7 cytotoxicity genes to determine the cytotoxicity score of γδ-T cells, CD8 T cells and NK cells, adapted from Tirosh, et al^59^. **c,** Score of cytotoxicity genes in Fig. 1b in NK, CD8 T and γδ-T cells. *Tukey’s multiple comparisons test for one-way ANOVA.* **d,** Heatmap representing the correlation between genes in specific cell subsets, which significantly interact with the indicated immune cell subset and significantly correlate with the cytotoxicity of said immune cell subset. Genes with at least one significant correlation, with either NK, CD8 T or γδ-T, were included. The black box indicates a negative sum of the correlations of the three immune cell subsets and represent the genes selected for further analysis. **e,** Correlation of 13 selected genes from Fig. 1d with cytotoxicity in dataset with bulk-RNA data from 498 neuroblastoma tumors (r2.amc.nl/; Tumor Neuroblastoma - SEQC - 498 - RPM - seqcnb1; GSE497104243). **f,** Survival analysis using the 13 selected genes from Fig. 1d using the SEQC bulkRNA cohort. Left panel represents event-free survival and right panel represents overall survival. **g,** Kaplan-Meier curve indicating event-free survival for high- or low expression of *MIF* (left panel, expression cutoff: 157.498) and *MDK* (right panel, expression cutoff: 216.718). **h,** Dotplot representing the expression of *MIF* and *MDK* in several cell subsets in scRNAseq dataset^30^.

To further shortlist which of these 13 genes could have the highest potential as a target for combinatorial immunotherapeutic approaches, we validated the correlation between their gene expression and the immune cell cytotoxicity score in two additional large, independent bulk-RNA sequencing datasets of 498 (SEQC) and 122 (COMBAT-Versteeg) neuroblastoma tumors^47,48^. Out of the 13 genes, Macrophage Migration Inhibitory Factor (*MIF)* and Midkine (*MDK)* negatively correlated with cytotoxicity in both datasets (Fig. 1e, Supplementary Fig. 1a-d). In addition, high *MIF* and *MDK* expression was significantly associated with inferior event-free and overall survival of neuroblastoma patients, which was independent of MYCN amplification (Fig. 1f, g, Supplementary Fig. 1e-g). The strong consistent correlation with poor cytotoxicity and poor survival led us to focus on MIF and MDK as prime targets to improve the durability of CAR T-cell therapy.

In our single cell RNA sequencing dataset, *MIF* and *MDK* were highly expressed by tumor cells. MIF was predicted to interact with CD8^+^ T cells through the CD74 receptor and MDK with NK cells through the SORL1 receptor (Fig. 1h and Supplement Fig. 1h). We confirmed the high expression of *MIF* and *MDK* by tumor cells in a published single-cell RNA sequencing dataset of 19 neuroblastoma tumors by Verhoeven *et al.*^49^(Supplement Fig. 1i). MIF and MDK are pleiotropic proteins with distinct and context-dependent functions including being described to be overexpressed in several cancers and linked to suppressive effects on T cell cytotoxicity^50–58^. Taken together, these data indicate a potentially immunosuppressive role of MIF and MDK in the neuroblastoma TME.

### MIF and MDK are abundantly secreted by neuroblastoma tumoroids

To further address the potential role for MIF and MDK in the modulation of tumor-infiltrating lymphocyte cytotoxicity, we confirmed their robust protein expression by mass-spectrometry of 11 neuroblastoma tumoroids. MIF was identified amongst the top 20 most abundant proteins in proteomics of whole tumoroid lysates (Supplement Fig. 2a). MDK was found in 8 of 11 tumoroids, albeit in lower abundance. However, importantly, considering that both MIF and MDK are secreted proteins, their intracellular level need not be high for them to have an extracellular immune-suppressive function.

Thus, to further assess whether these proteins may be altering lymphocyte function in the TME, we specifically quantified their secretion by neuroblastoma tumoroids. We selected three representative tumoroid models with distinct genetic backgrounds from which we collected the concentrated secreted proteins (secretome) for analysis by LC-MS (Fig 2a,b and Supplement Fig. 2b). Notably, both MIF and MDK were amongst the top 100 most abundantly secreted proteins in all three tumoroids (Fig. 2c). Taken together, these data firmly establish MIF and MDK as abundantly secreted immunosuppressive factors which may hamper cytotoxic function of T/NK cells in the TME.

**Figure 2:**
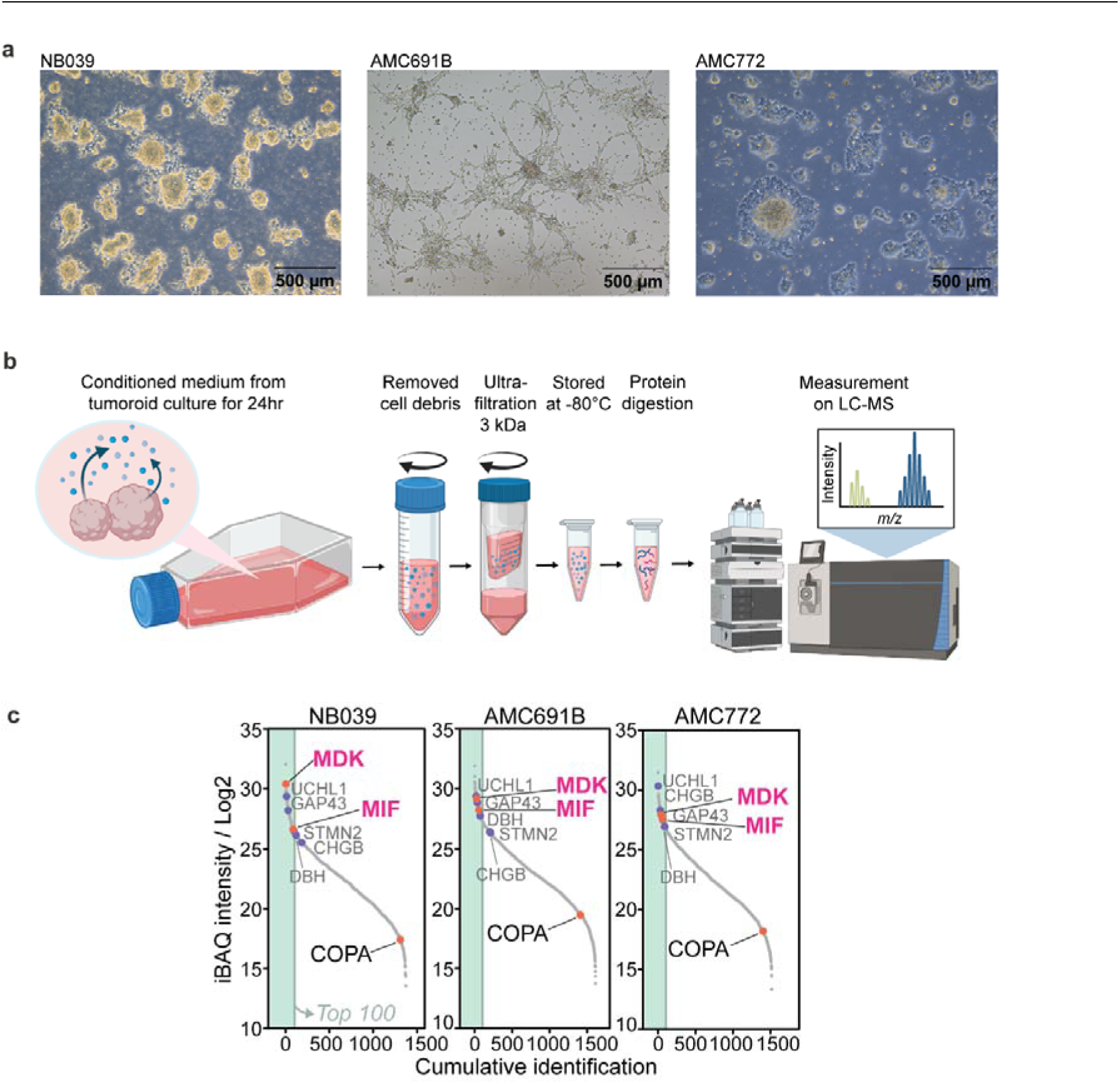
Identification of MIF and MDK in secretome of neuroblastoma tumoroids by LC-MS. **a**, Pictures of cultures from selected tumoroids for secretome collection. Scalebar indicates 500µM. **b,** Schematic overview of procedure to collect and process the secretome for analysis. Further details are provided in the methods section. Figure made with Biorender.com **c,** Liquid chromatography–mass spectrometry (LC-MS) analysis of conditioned medium from three neuroblastoma tumor organoids detailing their secretome. Red circles/black font: targets identified in figure 1d. Blue circles/grey font: neuroblastoma reference proteins. Left green bar indicates top 100 most abundant proteins. iBAQ (intensity Based Absolute Quantification) indicates the protein’s non-normalized intensity divided by the number of measurable peptides and, hence, indicating the relative abundance of the protein in the sample.

### MIF depletion increases CAR-T cell activation

Since CAR T-cell therapy is a promising upcoming treatment strategy for neuroblastoma, we further investigated the suppressive effect of MIF and MDK on T cells specifically. First, we activated healthy donor T cells with anti-CD3/CD28 stimulation in the presence and absence of recombinant MIF and MDK. Despite donor-variability, the presence of recombinant MIF and MDK indeed modestly decreased the activation of CD8+ T cells, as measured by expression of cytotoxicity marker Granzyme B, confirming our *in silico* findings (Fig. 3a, Supplement Fig. 3a). In addition, we observed a significant suppression of CD4+ and CD8+ T cell proliferation when co-incubated with MIF, while suppression by added MDK was not significant in CD8+ T cells (Fig. 3b, Supplement Fig 3b).

**Figure 3:**
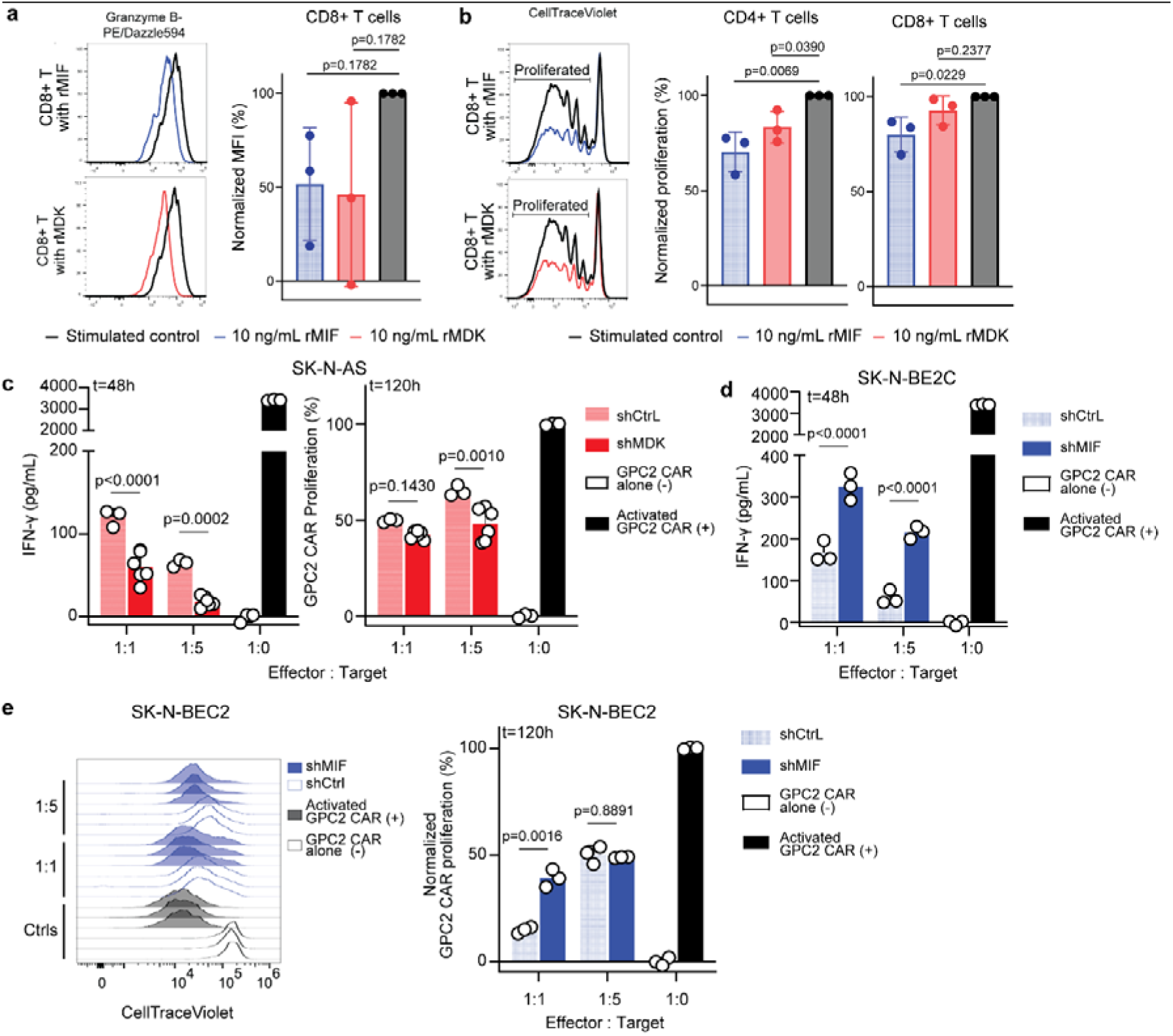
The immunosuppressive effect of MIF and MDK on CAR-T cell activation. **a**, Flow cytometry analysis of healthy donor peripheral blood T cells granzyme B MFI (Median Fluorescence Intensity) after 4 day *in vitro* stimulation with anti-CD3/anti-CD28 beads, in the presence of rMIF or rMDK (10 ng/mL). Representative graphs (left two panels) of granzyme B expression in CD8+ population. Combined data from three healthy donors (right panel). MFI was normalized to stimulated control. (*n=3 healthy donors,* blue indicated recombinant MIF, red indicates recombinant MDK). **b,** Proliferation analysis from same experiment as Fig. 3a. Representative graphs of CellTrace Violet peaks in CD8+ population (left panel). Combined data from three donors gated on CD4+ population (middle panel). Combined data from three donors gated on CD8+ population (right panel). Data is normalized to stimulated control (*n=3 healthy donors*). **c,** Activation of GPC2 CAR T-cells in co-culture with SK-N-AS models as measured by IFN-γ ELISA (left panel) and GPC2 CAR T-cell proliferation (right panel) in two effector : target ratios. SK-N-AS-shCtrl model (light red) and SK-N-AS-shMDK (dark red). Statistical analysis shows Two-way ANOVA with Šídák’s multiple comparisons test. (*n=1* CAR donor with several technical replicates). **d,** IFN-γ concentration in supernatant after 48 hours co-culture of GPC2 CAR T-cells with SK-N-BEC2-shMIF (dark blue) or shCtrl cells (light blue). Statistical test shows results for two-way ANOVA with Šídák’s multiple comparisons test, (*n=1* CAR donor with 3 technical replicates). **e,** Flow cytometry results from co-culture of GPC2 CAR-T cells with SK-N-BEC2-shMIF (dark blue) or shCtrl cells (light blue). Left panel shows proliferation after 5 days of co-culture for one representative CAR-T cell donor. Right panel normalized CAR T-cell proliferation, data were normalized to the activated control. Statistical test shows results for two-way ANOVA with Šídák’s multiple comparisons test.

We next used CAR T-cells targeting GPC2, which have shown robust safety and efficacy in diverse preclinical models of neuroblastoma and are currently being tested clinically (NCT05650749), to further investigate the roles of MIF and MDK in CAR T-cell modulation^44,60,61^. To study this interaction, we generated neuroblastoma cell lines with genetic depletion of MDK (SK-N-AS-shMDK) and MIF (SKNBE2C-shMIF) using shRNA-mediated silencing (Supplement Fig. 3c-g). Decreased MDK secretion unexpectedly decreased CAR T-cell activation and proliferation, indicating that it may not have the highest potential as therapeutic target (Fig. 3c). Interestingly, decreased MIF secretion did result in increased CAR T-cell activation in two out of three CAR T-cell donors, which was shown both by IFN-γ secretion and proliferation of CAR T-cells (Fig. 3d-e, Supplement Fig. 3h,i). Taken together, these mild, but significant effects support a clear functional role for MIF in modulating CAR T-cell activation in the neuroblastoma TME, while the role of MDK on T cell function remains unclear.

### Tumor-derived MIF reduces CAR-T cell cytotoxicity *in vitro* and *in vivo*

To further assess how MIF affects the function of CAR T cells, we next quantified the killing capacity of CAR T-cells against MIF-deficient neuroblastoma cells using co-incubation assays with SKNBE2C-shMIF and GPC2 CAR T-cells. *In vitro,* shMIF cells were killed significantly more efficiently by GPC2 CAR T-cells than control SKNBE2C cells (Fig. 4a,b). To validate these results *in vivo*, we generated SKNBE2C-shMIF or shCtrl neuroblastoma xenografts in NSG mice. We treated them, when flank tumors were established, with an intravenous dose of GPC2 CAR T-cells or CD19 CAR T-cells as negative control (Fig. 4c). GPC2 CAR-T cells killed MIF-deficient tumors more effectively than MIF-proficient tumors, leading to a significantly slower outgrowth of the tumors (Fig. 4d, Supplement Fig. 4a), and significantly prolonged survival (Fig. 4e). Intriguingly, MIF depletion by itself already reduced the growth rate of the tumors in vivo, possibly indicating an intrinsic dependency of neuroblastoma cells on MIF for proliferation and survival (Fig. 4d), which could make MIF a particularly promising target for therapeutic intervention. Taken together, these studies confirm that reduction of tumor-derived MIF secretion can enhance CAR-T cell efficacy both *in vitro* and *in vivo*.

**Figure 4:**
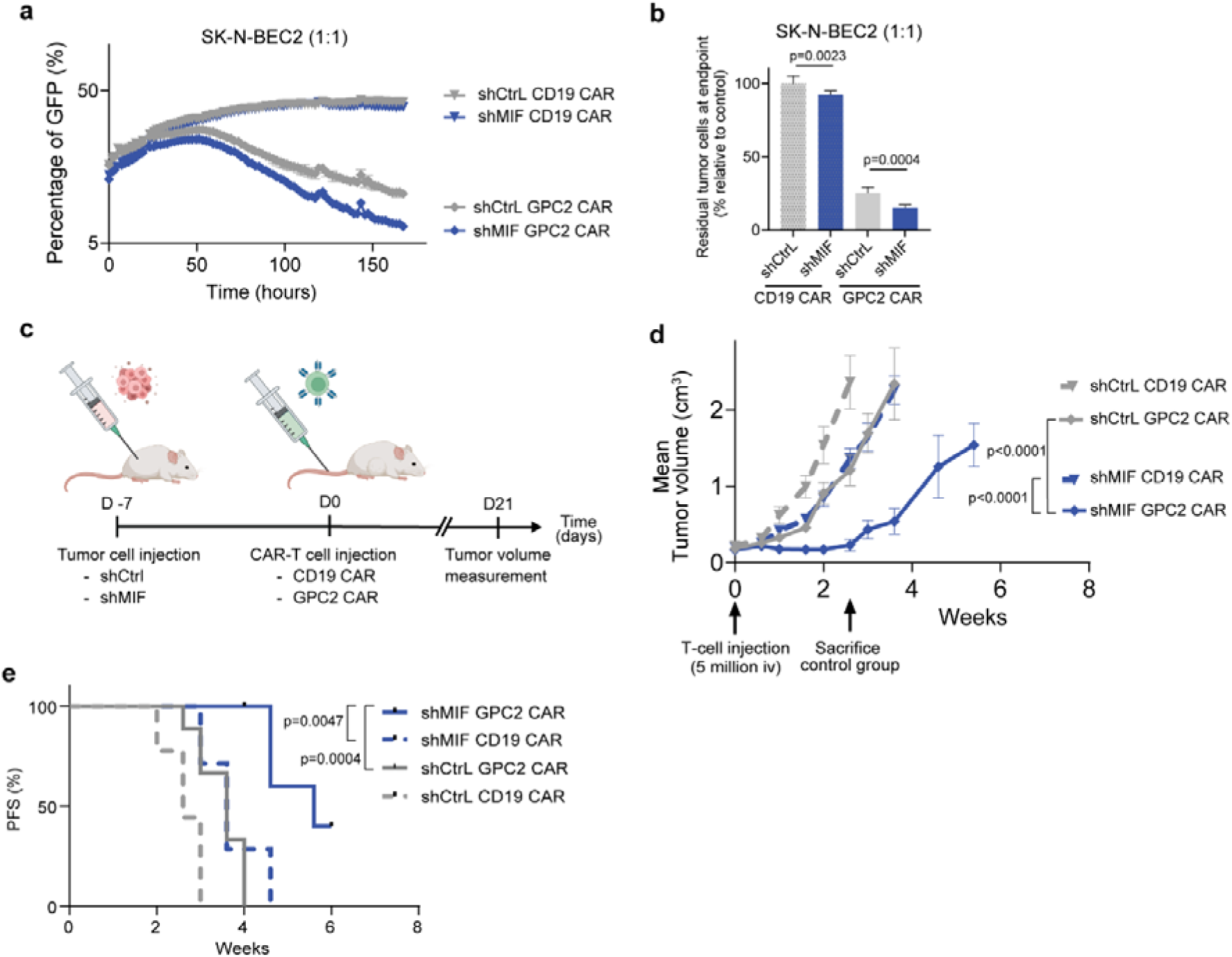
Reducing MIF secretion of the tumor increased the cytotoxicity of CAR-T cells. **a**, IncuCyte S3 experiment measuring tumor growth during co-culture with CAR-T cells. SK-N-BEC2 neuroblastoma cells with shCtrl (grey) or shMIF (blue) were culture with a control CAR-T cell targeting CD19 (triangles) or tumor antigen GPC2 (diamonds). Effector : Target ratio of 1:1. **b,** Normalized residual tumor cells at endpoint of the experiment in Fig 4a. One-way ANOVA statistical test with Holm-Šídák’s multiple comparisons test for significance. **c** Schematic overview of procedure to study the effect of MIF-knock down on cytotoxicity of CAR-T cells in mice. Figure made with Biorender.com. **d,** Average SK-N-BEC2 shCtrl or shMIF tumor growth. Measuring of tumor size started when CD19 or GPC2 CAR-T cells were injected (5×10^6^, iv, at arrow indication). Experimental groups of n=6 – n=9. Statistics show two-way ANOVA using Tukey’s multiple comparisons test at t=2.6 weeks, when mice in the control group were sacrificed. **e,** Progression free survival (PFS) of shCtrl or shMIF tumor bearing mice treated with CD19.CAR or GPC2.CAR. Statistical analysis shows p-values for a log-rank (Mantel-Cox) test.

### MIF degradation by PROTAC enhances neuroblastoma CAR T cell activation and cytotoxicity

To translate our findings towards a clinically applicable strategy, we explored the use of a MIF targeting PROteolysis Targeting Chimera (PROTAC) protein degrader. PROTAC constitutes a novel therapeutic modality using the target cell’s protein-degradation machinery to eliminate proteins of interest^62,63^. A PROTAC consist of a ligand for the protein of interest, a ligand for a E3 ligase and a linker that connects between the two ligands. Binding of the protein of interest in close proximity to the E3 ligase causes ubiquitination of the protein of interest, which initiates the degradation of this protein by the proteosome (Fig 5a)^62,63^.

**Figure 5:**
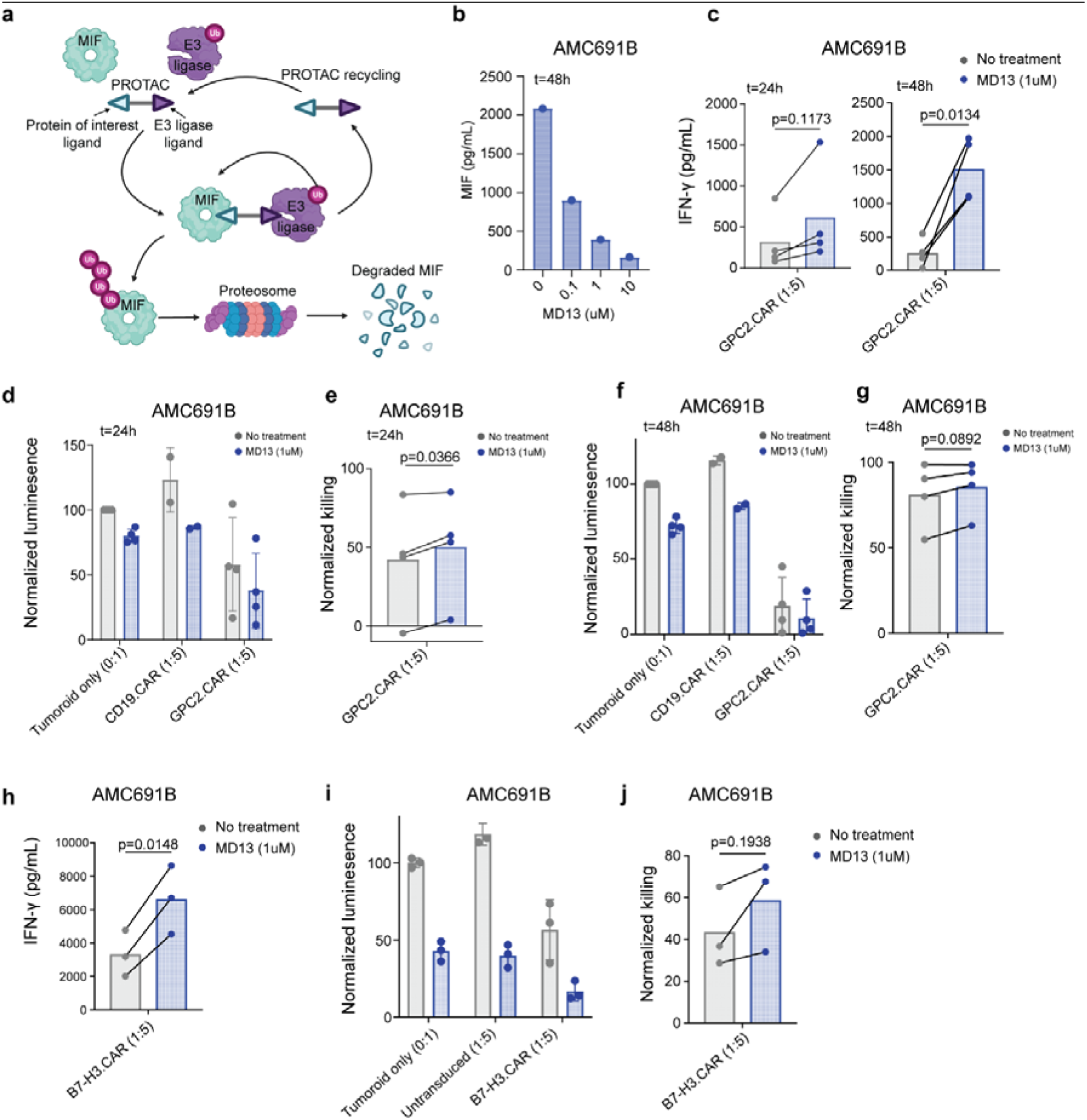
MIF PROTAC reduces MIF secretion, which allows for more efficient activation of CAR-T cells. **a**, Schematic representation of PROTAC degraders. Adapted from Dale, et al^63^. Figure made with Biorender.com **b**, MIF concentration AMC691B tumoroid model 48 hours of treatment with 0.1, 1 or 10µM MD13. Measured using Luminex. **c**, IFN-γ concentration of supernatant from co-culture of AMC691B without treatment (grey) or with 1uM MD13 treatment (blue) in combination with GPC2 CAR-T cell in a 1:5 Effector : Target ratio, as measured by ELISA. Left panel shows concentration at 24 hours and right panel shows concentration at 48 hours. Statistical analysis shows results for paired t-test. (*n=4* CAR T-cell donors) **d,** Luminescence signal of luciferase transduced tumoroid model AMC691B after co-culture of 24 hours. Normalized to untreated tumoroid only. Tumoroids were pre-treated for 48 hours before co-culture. (*n=2* CD19-CAR T-cell donors, *n=4* GPC2-CAR T-cell donors) **e,** Normalized GPC2 CAR-T cell killing from Fig 5d. Data were normalized to the tumoroid only untreated or treated control, respectively. Statistical analysis shows results for paired t-test. **f,** Luminescence signal of luciferase transduced tumoroid models after co-culture of 48 hours. Normalized to untreated tumoroid only. Tumoroids were pre-treated for 48 hours before co-culture. (*n=2* CD19-CAR T-cell donors, *n=4* GPC2-CAR T-cell donors) **g**, Normalized GPC2 CAR-T cell killing from Fig. 5f. Data were normalized to the tumoroid only untreated or treated control, respectively. Statistical analysis shows results for paired t-test. **h,** IFN-γ concentration of supernatant from co-culture of AMC691B without treatment (grey) or with 1µM MD13 treatment (blue) in combination with B7-H3 CAR-T cell in a 1:5 Effector : Target ratio, as measured by ELISA. (*n=3* B7-H3-CAR T-cell donors) **i,** Luminescence signal of luciferase transduced tumoroid models after co-culture of 24 hours. Normalized to untreated tumoroid only. Tumoroids were pre-treated for 48 hours before co-culture. (*n=3* CAR T-cell donors with untransduced controls). **j,** Normalized B7-H3 CAR T-cell killing from Fig. 5i. Data were normalized to the tumoroid only untreated or treated control, respectively. Statistical analysis shows results for paired t-test.

To assess whether MIF degradation by PROTAC would improve CAR T-cell efficacy, we treated a patient-derived tumoroid model with high MIF secretion (AMC691B) and a tumoroid model with lower MIF secretion (AMC691T) with MD13, a PROTAC targeting MIF^64^. Treatment with MD13 indeed reduced the concentration of secreted MIF in the culture supernatant of the neuroblastoma tumoroids, with a maximal reduction of 81-84% after 48 hours (Fig. 5b, Supplement Fig. 5a,b). We next combined MD13 treatment with CAR T-cells targeting GPC2, a target expressed by both tumoroid models (Supplement Fig. 5c). The tumoroid models were previously transduced with a constitutively active GFP/Luciferase construct for bio-luminescence based viability assays^65^. In co-cultures, tumoroids were pre-treated with MD13 for 48 hours to reduce MIF secretion before adding CAR T-cells. In line with our previous indications, GPC2 CAR T-cells proved significantly more activated after 48 hour co-cultures with MD13-treated AMC691B than untreated AMC691B (Fig. 5c).

This indicates that, indeed, MIF secreted by tumor cells may suppress CAR T-cell activation. Similar to our observation in the *in vivo* study, MD13 treatment alone reduced the viability of tumoroids by ∼20% after 24 hours (Fig. 5d). Addition of GPC2 CAR-T cells further reduced tumoroid viability to 37%, which, when normalized to the respective (treated or untreated) ‘tumoroid only’ controls, resulted in a significantly increased killing of MD13-treated tumoroids (Fig. 5d, e). After 48 hours, only a trend of increased killing was observed, since nearly all tumor cells were already killed even without MD13 treatment (Fig. 5f, g). The second tumoroid model (AMC691T), with lower endogenous MIF secretion, showed similar trends when treated with MD13 and GPC2 CAR T-cells, with significantly increased killing after 24 hours of co-culture (Supplement Fig. 5d-f).

To confirm the potential of MIF-degradation as an intervention to enhance CAR-T cell efficacy, we also tested the killing capacity and activation of B7-H3 targeting CAR T-cells when co-cultured with MD13-treated tumoroids. The antigen B7-H3 is also expressed by both tumoroid models (Supplement Fig. 5g). B7-H3 CAR T-cells were also significantly more activated against MD13-treated tumoroids, as measured by IFN-γ secretion, which resulted in increased killing of the tumoroids (Fig. 5h-j and Supplement Fig. 5h,I). These data confirm that MIF degradation results in increased activation of neuroblastoma-targeting CAR T-cells. Taken together, MIF PROTAC treatment provides a promising therapeutic strategy to reduce MIF secretion by neuroblastoma tumors and thereby enhance activation of clinically relevant CAR T-cells.

## Discussion

CAR T-cell therapy is a promising new approach for the treatment of cancer. However, in solid tumors, the immunosuppressive TME remains a major challenge for CAR T-cell persistence and durability. In this study, we identified MIF as a factor robustly secreted by neuroblastoma, which inhibits the optimal activation of CAR T-cells. Depletion of MIF genetically or via degradation by PROTAC treatment enhanced the efficacy of multiple neuroblastoma targeting CAR T-cells, together supporting the importance of this secreted protein in immune suppression.

We observed that neuroblastoma tumor cells express high levels of MIF and partly depend on MIF for their proliferation and/or survival, which is consistent with prior findings^66^. Interestingly, MIF was initially recognized as a lymphocyte-secreted factor influencing macrophage motility but was later described to be a multi-functional cytokine with context-dependent functions^50–52,67,68^. More recently, MIF has been described in the context of several malignancies, including neuroblastoma, to correlate with tumor growth, metastases and survival^54,55,69–71^. MIF upregulation in neuroblastoma can induce MYCN expression, resulting in tumor progression^72^. Moreover, MIF has been described to be involved in bone marrow metastases, where the expression of MIF contributes to survival and invasion of the tumor cells as well as an immunosuppressive tumor microenvironment^69,73^. Taken together, these studies suggest an important intrinsic role for MIF in neuroblastoma growth and progression, which makes MIF an attractive target for therapeutic intervention.

Next to these tumor-intrinsic effects, we found that tumor-secreted MIF reduces cytotoxicity of immune cells in the tumor microenvironment of neuroblastoma. This has also been described in other malignancies. In soft-tissue sarcoma, tumor-secreted MIF can shape macrophages to become more pro-tumorigenic and in melanoma, MIF has been described to tune myeloid-derived suppressor cells to hinder cytotoxicity of T cells^74,75^. In rhabdomyosarcoma, MIF knock-out increased the potential of cellular immune interventions^76^. In addition, a recent study presenting resistance mechanisms of BCMA CAR T-cell therapy in multiple myeloma treatment, revealed MIF as one of the immunosuppressive signaling pathways^77^. These studies corroborate our findings of an immunosuppressive effect of MIF on CAR T-cells targeting neuroblastoma.

While the direct effect of MIF on T cells has been scarcely studied, in our single cell RNA sequencing analysis MIF was predicted to interact directly with T cells through the CD74 receptor. CD74 is the invariant chain of major histocompatibility complex II (MHC-II). Extracellular binding of MIF induces heterodimerization with CD44 and subsequent ERK-MAPK pathway activation, resulting in, amongst others, prostaglandin E2 (PGE2) production, a well-known suppressor of T cell activation^53,78^. Others showed that MIF can cause activation-induced T cell death through the IFN-γ pathway^79^. The exact mechanism through which MIF directly inhibits T cells still remains to be further unraveled. However, several studies describe the effect of MIF on other cells in the TME, e.g. myeloid-derived suppressor cells or M2-like macrophages, which could indirectly affect T cells ^80,81^. This highlights the importance of studying the immunomodulatory role of MIF in immunocompetent *in vivo* models.

Since MIF is a secreted factor, it may exert immunosuppressive effects beyond the local tumor microenvironment and affect CAR T-cells systemically. In gastric cancer, head and neck squamous cell carcinoma and hepatocellular carcinoma patients, MIF levels were elevated in the plasma from patients with accelerated cancer progression and plasma-MIF concentration correlated negatively with response to immune-checkpoint inhibition^82–85^. This suggests MIF in patient serum could not only be used as a biomarker for more severe disease, but, by causing an immunosuppressive systemic environment, it may also impact CAR T-cells systemically, reducing their cytotoxic potential and possibly their ability to migrate towards and into the tumor. Patient-specific assessment of MIF levels could therefore even be considered as a biomarker for patient stratification, to tailor treatment approaches.

A unique aspect of our study is our multi-omics approach, as secretion, *per se*, of immuno-suppressive factors does not necessarily correlate with RNA expression. By combining two sets of unsupervised analyses, i.e. single-cell RNA sequencing and secretome proteomics, we prioritized MIF and MDK as candidate targets. Moreover, we were able to validate *in silico* findings *in vitro* and *in vivo* for MIF using clinically-relevant CAR T-cells. Furthermore, we degraded MIF with PROTAC treatment, providing a potential therapeutic intervention to improve CAR T-cell efficacy. However, the potential clinical use of MD13 or related PROTACs will have to be validated further to evaluate its safety profile and efficacy *in vivo*.

Fundamentally, our study reveals critical insights into the immunosuppressive tumor microenvironment of neuroblastoma. This has an important impact on the development and application of immunotherapies for neuroblastoma, especially for adoptive cell therapy. By understanding how neuroblastoma suppresses the activation of CAR T-cells and their migration to the tumor, we can further optimize these therapies for clinical application. This could include treating patients with a MIF-PROTAC before administering CAR T-cell therapy to ensure optimal efficacy of the CAR T-cell product. The MIF-PROTAC tested in this study, MD13, still needs further investigation before it could be clinically applied. However, other interventions against MIF are in clinical development. For example, several monoclonal antibodies targeting MIF showed promising results *in vitro* and *in vivo,* and led to a first clinical study in patients with solid tumors (clinicaltrials.gov; NCT01765790). The results of this study are still pending^86^. Based on the broad expression of MIF across solid tumors, inhibition or degradation of MIF may be a viable strategy to improve CAR T-cell efficacy not only in neuroblastoma but in many solid cancers. Moreover, our multi-omics pipeline to identify targetable immunosuppressive factors may also be applicable to other difficult to treat solid cancers to improve immunotherapy efficacy. In conclusion, we showed that selective targeting of immunosuppressive secreted factors in the TME, and MIF in particular, may improve the efficacy of CAR T-cell therapy for patients with solid tumors and may improve their treatment outcome.

## Methods

### Ethics statement

This study complies with all relevant ethical regulations. Studies involving primary tumor or patient data, were performed with data published before according to ethical approval. Animal experiments were conducted using protocols approved by the CHOP Institutional Animal Care and Use Committee (IACUC; Protocol #1464) with adherence to the NIH guide for the Care and Use of Laboratory Animals accredited by the Association for Assessment and Accreditation of Laboratory Animal Care (AAALAC).

### Transcriptomics analyses

Datasets used for transcriptomics analyses were previously published by our and other groups^30,47,49,87^. Cell isolation, library preparation, and cluster annotations are described in the publications. Correlation between cell subsets was predicted using the CellChat algorithm to perform an unbiased ligand-receptor analysis^45^. Predicted interactions were overlaid with the Human Protein Atlas database to select for genes which are either expressed on the surface membrane or secreted. Functionally relevant correlations were selected by correlating the predicted interaction with cytotoxicity of T/NK cell subsets based on an adapted cytotoxicity score from Tirosh, et al^59^. Only the interactions with a significant correlation with the cytotoxicity score (p<0.05) and the interactions which were significantly predicted to have an interaction between the cell subset X and T/NK cell subset Y (p<0.05) are shown.

### Tumoroid cultures

Patient derived tumoroids were previously established as described^65,88,89^. Tumoroids were cultured in optimized medium (DMEM, low glucose, GlutaMAX supplemented with 20% F-12 Ham’s Nutrient Mix, 100U/mL penicillin, 100ug/mL streptomycin, B27 (50x), N2 (100x), hIGF (300ng/mL), hFGF (40ng/mL), hEGF (20ng/mL), PDGFaa (10ng/mL), PDGFbb (10ng/mL)). Cells were passaged 1-2 times a week by breaking down larger spheres mechanically and moving the cells to a larger flask while refreshing the culture medium. Cells were propagated at 37°C in 5% CO_2_. For killing assays, tumoroids transduced with a GFP-Luciferase construct were used to be able to measure viability of the tumor cells^65,88^.

### Tumoroid collection and whole proteomics

For whole proteomics of tumoroid models, the tumoroid lines were expanded in culture as described above to one full T175 flask. Cells are harvested by pipetting the cells loose from the flask or with enzymatic detachment using TrypLE (ThermoFisher) or Accutase (Sigma Aldrich). Cells are washed by spinning the Falcon tube with the harvest at 300g for 5 minutes and resuspending the pellet in PBS. After washing twice, the pellet is left to dry and stored at -80°C until further sample preparation. Sample preparation and mass spectrometry were performed as published before^90^.

### Tumoroid secretome collection

Tumoroid cultures were expanded to two full T175 flasks. Medium was washed off and 45 mL empty DMEM GlutaMAX was left on for 24 hours to condition. The conditioned medium was spun down 5 minutes at 250g to get rid of any cells, after which the medium was spun down twice at 3220g for 10 minutes to get rid of any debris. The conditioned medium was concentrated by filtering through a 3kDa molecular weight cut-off Millipore filter down to 1mL. Proteins were denatured in 4M Urea, in a Tris-buffered environment at pH 8.0. Secreted proteins were reduced in 10mM Dithiothreitol (DTT; Sigma-Aldrich) at 20°C for 1 hour, and then alkylated in 20mM iodoacetamide (IAA; Sigma-Aldrich) at 20°C for 0.5 hour in the dark. Protein digestion was performed sequentially with Lys C (1:50) and Trypsin (1:50) for 4 and 16 hours respectively. Digested peptides were acidified to 2% formic acid and diluted to 1 mL for peptide cleanup by Sep-Pak C18 1cc (Waters). Desalted peptides were dried by vacuum centrifugation, and stored at -80°C for further use.

### Secretome sample preparation and LC-MS

Peptides were reconstituted with 2% formic acid and analyzed in triplicates on an Orbitrap Exploris 480 mass spectrometer (Thermo Scientific) coupled to an UltiMate 3000 UHPLC system (Thermo Scientific), that consisted of an µ-precolumn (C18 PepMap100, 5 µm, 100 Å, 5 mm × 300 µm; Thermo Scientific), and an analytical column (120 EC-C18, 2.7 µm, 50 cm × 75 µm; Agilent Poroshell). Solvent A was made of 0.1% formic acid in water and solvent B was 0.1% formic acid in 80% acetonitrile, 20% water. Peptides were resolved on a 175min gradient from 10 to 40% Solvent B. Mass spectrometry data were acquired in data-dependent acquisition mode. MS1 scans were acquired between m/z 375-1600 at a resolution of 60,000, upon signal accumulation to AGC target of 1×10^6^. Multiply charged precursors starting from m/z 120 were selected for further fragmentation. Higher energy collision dissociation (HCD) was performed with 28% normalized collision energy (NCE), at a resolution of 30,000, upon signal accumulation to AGC target of 1e5. An isolation window of 1.4 m/z and dynamic exclusion of 24s were used.

### Secretome data analysis

MS raw files were searched with MaxQuant (version 1.6.10.0) against the human UniProt database (version April 22, 2021) using the integrated Andromeda search engines. Protein N-terminal acetylation and methionine oxidation were added to variable modification, whereas cysteine carbamidomethylation was added to fixed modification. Trypsin/P was set as the enzyme for digestion and up to 2 miss cleavage was allowed. Precursor ion tolerance was set to 20_Jppm for the first search and 4.5_Jppm after recalibration, and fragment ions tolerance was set to 20_Jppm. False discovery rate (FDR) of 1% was set at both peptide spectrum match (PSM) and protein level by using a reverse decoy database strategy. Label-free quantification (LFQ) algorithm and the match-between-run feature were enabled for protein identification. Proteins identified from empty DMEM GlutaMAX were excluded from the identifications.

### PBMC isolation

Peripheral blood mononuclear cells (PBMCs) for functional assays such as suppression assays and CAR-T cell production were isolated from healthy donor blood. Whole blood samples were centrifuged at 350g for 10 minutes at room temperature with the brake at 0, after which the plasma could be taken off. The concentrated blood was diluted with basic RPMI medium (supplemented with 2% FBS). Lymphocytes were further isolated from the blood using Ficoll-Paque (Sigma Aldrich) density centrifugation, after which the buffy coat layer could be pipetted off. Isolated cells were washed twice, counted and stored in liquid nitrogen storage. Primary human T-cells were obtained from anonymous donors through the Human Immunology Core at the University of Pennsylvania under a protocol approved by the CHOP Institutional Review Board. Donors provided informed consent through the University of Pennsylvania Immunology Core.

### Suppression assays

Healthy donor PBMC were stained with anti-CD3-AF700 (300324, Biolegend, 1:400) and fixable viability dye efluor 506 (Invitrogen, cat. 65-0866-14) for FACS sorting of live CD3+ T cells on a Sony SH800S machine. Sorted T cells were labelled with 2μM CellTrace Violet (Invitrogen, cat. C34557) and stimulated with α-CD3/α-CD28 Dynabeads (Gibco, cat. 11131D) in the presence of 10 ng/mL recombinant MIF (Biolegend, cat. 599404) or MDK (Biolegend, cat. 754104) for 4 days. After 4 days, cells were washed and stained for flow cytometry analysis.

### Cell lines and modifications

Neuroblastoma cell lines SY5Y, SK-N-AS, SKNBE2C and Kelly were cultured in RPMI 1640 with 10% FBS, 1% L-Glutamine and 1% Penicillin-streptomycin at 37°C with 5% CO_2._ Cells were passaged twice a week and routinely authenticated. Short-hairpin RNA (shRNA) constructs were obtained from Sigma (TRCN0000303918, TRCN0000331210, TRCN0000331252, TRCN0000331270, TRCN0000331211, TRCN0000056818, TRCN0000056819, TRCN0000056820, TRCN0000056821, TRCN0000056822) and used to prepare MIF and MDK knock downs, using lentiviral transduction of several shRNA construct.

### Western Blot

Protein lysates from neuroblastoma cells were separated on 4-12% Bis-Tris gels (Life Technologies), transferred to a PVDF membrane, blocked in 5% non-fat milk in Tris-buffered saline and Tween-20 (TBS-T), and blotted using standard protocols. Membranes were incubated at 4°C overnight in MIF (Cell Signaling; #87501; 1:1000), MDK (Santa Cruz; #46701; 1:200) or β-actin antibodies, washed x 3 in TBS-T and developed with a chemiluminescent reagent (SuperSignal West Femto, Thermo Fisher Scientific).

### ELISA

For determining the concentration of MIF and MDK secreted by the mutants, we used Human MIF and midkine DuoSet ELISA kits (DY289 and DY258, respectively; R&D systems). Cells were plated in a 96-well plate and supernatant was collected at several time point. To account for the differences in number of cells, the luminescence signal by the GFP-luciferase construct was measured. IFN-γ secretion to validate CAR-T cell activation was measured using the Human INF-γ ELISA kit (Biolegend) or Human IFN-γ DuoSet ELISA kit (R&D Systems). Supernatant of co-cultures is harvested at indicated times and stored at -20°C until analyzed.

### Flow Cytometry

Single cell suspensions were incubated with indicated antibodies and run on a Beckman CytoFLEX S or LX cytometer (Beckam Coulter). CD3-AF700 (300324, Biolegend, 1:400), CD4-FITC (357406, Biolegend, 1:400), CD8-PerCP (344707, Biolegend, 1:100) and Granzyme-B-PE/Dazzle 594 (372215, Biolegend, 1:100) were used to stain healthy donor T cells in suppression assays. For intracellular staining of Granzyme-B, the cells were fixated using Fix/Perm kit (Invitrogen; FOXP3). CD45-APC, PE or FITC (J33, Beckman Coulter; 1:10) or CD3-APC, PE or FITC (UCHT1; Beckman Coulter; 1:10) were used to stain T-cells in co-culture assays. GPC2 CAR expression was determined using APC or PE-conjugated human recombinant GPC2 protein (R&D Systems; 2304-GP-050; 1:500) and CD19 CAR using PE-tagged protein L (58036S; Cell Signaling). B7-H3 CAR expression was determined using CD34-APC (FAB7227R, QBend10 clone, R&D, 1:100), which binds to the co-expressed marker RQR8. For in vitro T-cell proliferation, we stained cells with CellTrace™ CFSE or Violet Cell Proliferation Kit (C34554 and C34571, respectively; ThermoFisher). Data were analyzed using FlowJo software.

### GPC2 CAR-T cell production

CAR constructs were generated using a lentiviral construct backbone containing an EF-1α promoter. GPC2.CARs were designed using a CD8α leader, followed by the single-chain variable fragment (scFv) of the GPC2 D3 Barisa antibody with a VL-VH orientation with (Gly4Ser)x3 linker, a CD28 hinge and transmembrane domain, a 41BB and CD3-ζ co-stimulatory domains. CD19.CAR was designed using a CD8α leader, followed by a scFv derived from the FMC63 antibody, a CD8 hinge and transmembrane domain, and a 4-1BB and CD3-ζ co-stimulatory domains. DNA transfections, lentivirus production using second and third-generation lentiviral systems and virus transductions were performed as previously described^61^.

Primary T-cells (CD4 and CD8; 1:1 ratio) were activated for 24 hours with Dynabeads™ Human T-Expander CD3/CD28 (Thermo Fisher) at 3:1 bead: T-cell ratio together with human recombinant IL-15 and IL-7 (PeproTech; 5 ng/mL) in AIM-V medium supplemented with 5% FBS, 2mM L-Glutamine, 0.1M HEPES buffer and 1% streptomycin/penicillin at a density of 1×10^6^ cells per mL. On day 2, T-cells were transduced with CAR-containing lentiviral particles and maintained in culture until days 5-7. Then, beads were magnetically removed, and primary T-cells were placed in vented Erlenmeyer flasks at a density of 0.25×10^6^ cells per mL and cultured in agitation (125 rpm) until day 12-15. At that timepoints, cells were collected, cell viability and CAR expression determined, and cells frozen until functional in vitro or in vivo assays.

### B7-H3 CAR-T cell production

For the production of B7-H3 targeting CAR-T cells, we used previously published TE9-28z CAR-T cell^43^. Phoenix-ampho cells, transduced to constitutively express the CAR construct were plated with 5×10^6^ cells in a T175. After 72 hour incubation, the retroviral supernatant was collected and spun down at 500g for 5 minutes to get rid of any cells and debris. Aliquots of 5mL virus were snap frozen in liquid nitrogen and stored at -70°C. Healthy donor PBMCs were either used fresh from isolation or from frozen vials. PBMCs were activated with α-CD28 (130-093-375, Miltenyi, 0.5µg/mL) and α-CD3 (130-093-387, Miltenyi, 0.5µg/mL) for 24 hours, after which 100 iU/mL IL-2 (130-097-748, Miltenyi) was added and left to incubate for another 24 hours. Non-tissue culture treated 24-well plated were coated overnight with 8µL/mL RetroNection (T100A, Tekada). 3×10^5^ activated PBMCs were plated per well with 100 iU/mL IL-2 and 1.5 mL thawed or fresh retrovirus was added. The plates were spun for 40 minutes at 32°C with brake at 0, after which the plates were incubated at 37°C with 5% CO_2_ for 72 hours. The virus was washed of and CAR-T cells were expanded by refreshing the medium every two days to make a dilution of 1×10^6^ cells per mL and fresh IL-2 (100 iU/mL) was added. CAR expression was determined on day 3 by flow cytometry.

### Killing assays *in vitro*, using IncuCyte readout

CAR T-cell killing was evaluated using IncuCyte-based assays. GFP-positive, MIF wild-type or knock-down neuroblastoma cells were cultured in 96-well plates and co-incubated with CD19 or GPC2.41BBz CAR T-cells at a 1:1 effector:target (E:T) ratio. Plates were imaged every 1-2 hours using the IncuCyte ZOOM Live-Cell analysis system (Essen Bioscience). Total integrated GFP intensity per well was assessed as a quantitative measure of viable tumor cells. Values were normalized to the starting measurement and plotted over time.

### Killing assay *in vitro*, using luminescence readout

Tumoroid transduced with a GFP-luciferase construct were cultured as described above. A single cell suspension was prepared with Accutase (Sigma Aldrich) and mechanical dissociation of the tumoroids. 10.000 single cells were plated and rested for 2-3 days to reform spheres. Effector cells were added at t=0 and left to incubate until indicated timepoints. Supernatants were collected for ELISA and D-luciferin (122799, PerkinElmer, 150ug/mL) was added to the wells and incubated for 5 minutes at 37°C. Luminescence signal was measured with the FLUOstar Omega microplate reader. All assays were performed with three technical replicates.

### Mouse study

To generate SKNBE2C WT or MIF knock-down xenografts, a total of 5×10^6^ SKNBE2C cells were injected subcutaneously in subcutaneously in the flanks of 6-week-old immunodeficient female NSG mice (NOD-scid IL2Rgammanull; 005557; Jackson Labs) using 100 µL of Matrigel (Corning). Once tumors reached a volume of 0.15-0.3 cm^3^, a total of 5×10^6^ CAR+ T-cells were administered intravenously in 100uL of PBS. Mouse weights and tumor volumes were measured at least twice weekly and tumor volumes were calculated as volume =((diameter 1/2+diameter 2/2)⁄(3*0.5236))/100. Mice, unless otherwise noted, were treatment naive and maintained in cages of up to 5 mice under barrier conditions with ready access to feed and water following IACUC guidelines (Protocol #643).

### Statistics and reproducibility

Statistical tests were performed using GraphPad Prism v10.0.2 software. Which test is used is specified in the figure legends. A p-value of <0.05 was considered statistically significant, unless stated otherwise. In some graphs, p<0.0001 was indicated, as GraphPad Prism does not provide the exact values of p when lower than 0.0001.

## Competing interests

K.R.B. and G.P.P have applied for patents for the discovery and development of immunotherapies for cancer, including patents related to GPC2-directed immunotherapies.

K.R.B receives royalties from Tmunity/Kite Pharma and ConjugateBio, Inc. for licensing of GPC2-related technology and funding from Tmunity/Kite Pharma and ConjugateBio, Inc. for research on GPC2-directed immunotherapies. K.R.B. is on the ConjugateBio Scientific Advisory Board. M.B. holds patents pertinent to cellular immunotherapy development and manufacture, and has consulted for Lava Therapeutics. J.A. holds founder stock in Autolus ltd, consults for Roche and BMS, and holds patents in CAR-T design. J.M. has received research funding from Roche for in vitro work. No other conflicts of interest are declared.

## Acknowledgements

This work received funding from Villa Joep, Veni (Release the beast: Boosting CAR-T cell immunotherapy for neuroblastoma), Alex’s Lemonade Stand Foundation (G.P.P), NCI R37 CA282041 (KRB), NCI K08 CA230223 (KRB), and ProCan® and J.M. are supported by Australian European Union grant (GNT1170739, a companion grant to support the European Commission’s Horizon 2020 Program, H2020-SC1-DTH-2018-1, ’iPC - individualizedPaediatricCure’ [ref. 826121]). J.S. received travel grants from the Prins Bernhard Cultuurfonds and the KNAW Ter Meulen grant for this project. T.M. and L.V. (single cell genomics facility of Princess Máxima Center for Pediatric Oncology) are funded by KiKa.

W.W. is supported by Singapore Immunology Network (SIgN), Agency for Science, Technology and Research (A∗STAR); Biomedical Research Council (BMRC) Core Research Fund for use-inspired basic research (UIBR) and IAF-PP project H22J2a0043, and Singapore National Medical Research Council (NMRC) project MOH-001401-00.

## Data availability

Single-cell RNA sequencing data for analyses in figure 1 were published before by our group (DOI: https://doi.org/10.1016/j.ccell.2023.12.008). Proteomics data for figure 2c and supplementary figure 2a are available on Proteomics IDEntifications Database (PRIDE) (PXD051044).

## References

1. Galluzzi, L., Chan, T. A., Kroemer, G., Wolchok, J. D. & López-Soto, A. The Hallmarks of Successful Anticancer Immunotherapy. Sci. Transl. *Med* vol. 10 http://stm.sciencemag.org/ (2018).

2. June, C. H., O’Connor, R. S., Kawalekar, O. U., Ghassemi, S. & Milone, M. C. CAR T cell immunotherapy for human cancer. Science (1979) 359, 1361–1365 (2018).

3. Park, J. H. et al. Long-Term Follow-up of CD19 CAR Therapy in Acute Lymphoblastic Leukemia. New England Journal of Medicine 378, 449–459 (2018).

4. Lee, D. W. et al. T cells expressing CD19 chimeric antigen receptors for acute lymphoblastic leukaemia in children and young adults: A phase 1 dose-escalation trial. The Lancet 385, 517–528 (2015).

5. Mohty, M. et al. CD19 chimeric antigen receptor-T cells in B-cell leukemia and lymphoma: current status and perspectives. Leukemia 33, 2767–2778 (2019).

6. Zappa, E. et al. Adoptive cell therapy in paediatric extracranial solid tumours: current approaches and future challenges. Eur J Cancer 194, 113347 (2023).

7. Del Bufalo, F. et al. GD2-CART01 for Relapsed or Refractory High-Risk Neuroblastoma. N Engl J Med 388, 1284–1295 (2023).

8. Chavez, J. C., Bachmeier, C. & Kharfan-Dabaja, M. A. CAR T-cell therapy for B-cell lymphomas: clinical trial results of available products. Ther Adv Hematol 10, 204062071984158 (2019).

9. Matthay, K. K. et al. Neuroblastoma. Nat Rev Dis Primers 2, 16078 (2016).

10. Maris, J. M., Hogarty, M. D., Bagatell, R. & Cohn, S. L. Neuroblastoma. Lancet 369, 2106–2120 (2007).

11. Seeger, R. C. et al. Association of Multiple Copies of the N-myc Oncogene with Rapid Progression of Neuroblastomas . New England Journal of Medicine 313, 1111–1116 (1985).

12. Irwin, M. S. et al. Revised Neuroblastoma Risk Classification System: A Report From the Children’s Oncology Group. Journal of Clinical Oncology JCO.21.00278 (2021) doi:10.1200/JCO.21.00278.

13. Yu, A. L. et al. Anti-GD2 Antibody with GM-CSF, Interleukin-2, and Isotretinoin for Neuroblastoma. New England Journal of Medicine 363, 1324–1334 (2010).

14. Yu, A. L. et al. Long-term follow-up of a phase III study of ch14.18 (dinutuximab) + cytokine immunotherapy in children with high-risk neuroblastoma: COG study ANBL0032. Clinical Cancer Research 27, 2179–2189 (2021).

15. Yu, L. et al. GD2-specific chimeric antigen receptor-modified T cells for the treatment of refractory and/or recurrent neuroblastoma in pediatric patients. J Cancer Res Clin Oncol 148, 2643–2652 (2022).

16. Park, J. R. et al. Adoptive Transfer of Chimeric Antigen Receptor Re-directed Cytolytic T Lymphocyte Clones in Patients with Neuroblastoma. Molecular Therapy 15, 825– 833 (2007).

17. Straathof, K., et al. Abstract CT145: A Cancer Research UK phase I trial of anti-GD2 chimeric antigen receptor (CAR) transduced T-cells (1RG-CART) in patients with relapsed or refractory neuroblastoma. Cancer Res 78, CT145–CT145 (2018).

18. Louis, C. U. et al. Antitumor activity and long-term fate of chimeric antigen receptor– positive T cells in patients with neuroblastoma. Blood 118, 6050–6056 (2011).

19. Yu, L. et al. GD2-specific chimeric antigen receptor-modified T cells for the treatment of refractory and/or recurrent neuroblastoma in pediatric patients. J Cancer Res Clin Oncol 148, 2643–2652 (2022).

20. Dagher, O. K. & Posey, A. D. Forks in the road for CAR T and CAR NK cell cancer therapies. Nat Immunol 24, 1994–2007 (2023).

21. Dagar, G. et al. Harnessing the potential of CAR - T cell therapyL: progress , challenges , and future directions in hematological and solid tumor treatments. J Transl Med 3, 1–36 (2023).

22. Marofi, F. et al. CAR T cells in solid tumors: challenges and opportunities. Stem Cell Res Ther 12, 81 (2021).

23. Richards, R. M., Sotillo, E. & Majzner, R. G. CAR T Cell Therapy for Neuroblastoma. Front Immunol 9, 2380 (2018).

24. Wienke, J. et al. The immune landscape of neuroblastoma: Challenges and opportunities for novel therapeutic strategies in pediatric oncology. Eur J Cancer 144, 123–150 (2021).

25. Rahbar, M., Mehrazma, M. & Karimian, M. Tumor infiltrating cytotoxic CD8 T-Cells predict clinical outcome of neuroblastoma in children. Indian Journal of Medical and Paediatric Oncology 39, 159–164 (2018).

26. Mina, M. et al. Tumor-infiltrating T lymphocytes improve clinical outcome of therapy-resistant neuroblastoma. Oncoimmunology 4, e1019981 (2015).

27. Majzner, R. G. et al. Assessment of programmed death-ligand 1 expression and tumor-associated immune cells in pediatric cancer tissues. Cancer 123, 3807–3815 (2017).

28. Melaiu, O. et al. PD-L1 Is a Therapeutic Target of the Bromodomain Inhibitor JQ1 and, Combined with HLA Class I, a Promising Prognostic Biomarker in Neuroblastoma. Clin Cancer Res 23, 4462–4472 (2017).

29. Xin, C. et al. CD200 is overexpressed in neuroblastoma and regulates tumor immune microenvironment. Cancer Immunology, Immunotherapy 69, 2333–2343 (2020).

30. Wienke, J. et al. Integrative analysis of neuroblastoma by single-cell RNA sequencing identifies the NECTIN2-TIGIT axis as a target for immunotherapy. Cancer Cell (2024) doi:10.1016/j.ccell.2023.12.008.

31. Floutsis, G., Ulsh, L. & Ladisch, S. Immunosuppressive activity of human neuroblastoma tumor gangliosides. Int J Cancer 43, 6–9 (1989).

32. Li, R., Gage, D., McKallip, R. & Ladisch, S. Structural characterization and in vivo immunosuppressive activity of neuroblastoma GD2. Glycoconj J 13, 385–9 (1996).

33. Ladisch, S. et al. Shedding of GD2 ganglioside by human neuroblastoma. Int J Cancer 39, 73–6 (1987).

34. Burga, R. A. et al. Engineering the TGFb receptor to enhance the therapeutic potential of natural killer cells as an immunotherapy for neuroblastoma. Clinical Cancer Research 25, 4400–4412 (2019).

35. Soldati, R. et al. Neuroblastoma triggers an immunoevasive program involving galectin-1-dependent modulation of T cell and dendritic cell compartments. Int J Cancer 131, 1131–1141 (2012).

36. Mussai, F. et al. Neuroblastoma Arginase Activity Creates an Immunosuppressive Microenvironment That Impairs Autologous and Engineered Immunity. Cancer Res 75, 3043–3053 (2015).

37. Quinn, J. J. & Altman, A. J. The multiple hematologic manifestations of neuroblastoma. Am J Pediatr Hematol Oncol 1, 201–5 (1979).

38. Scott, J. P. & Morgan, E. Coagulopathy of disseminated neuroblastoma. J Pediatr 103, 219–22 (1983).

39. Evans, A. E. et al. Factors influencing survival of children with nonmetastatic neuroblastoma. Cancer 38, 661–6 (1976).

40. Chung, H. S., Higgins, G. R., Siegel, S. E. & Seeger, R. C. Abnormalities of the immune system in children with neuroblastoma related to the neoplasm and chemotherapy. J Pediatr 90, 548–554 (1977).

41. Wang, X. et al. Diminished cytolytic activity of γδ T cells with reduced DNAM-1 expression in neuroblastoma patients. Clinical Immunology 203, 63–71 (2019).

42. Szanto, C. L. et al. Immune Monitoring during Therapy Reveals Activitory and Regulatory Immune Responses in High-Risk Neuroblastoma. Cancers (Basel) 13, 2096 (2021).

43. Birley, K. et al. A novel anti-B7-H3 chimeric antigen receptor from a single-chain antibody library for immunotherapy of solid cancers. Mol Ther Oncolytics 26, 429–443 (2022).

44. Heitzeneder, S. et al. GPC2-CAR T cells tuned for low antigen density mediate potent activity against neuroblastoma without toxicity. Cancer Cell 40, 53–69.e9 (2022).

45. Jin, S. et al. Inference and analysis of cell-cell communication using CellChat. Nat Commun 12, (2021).

46. Wang, R. et al. SEPDB: a database of secreted proteins. Database 2024, (2023).

47. SEQC/MAQC-III Consortium. A comprehensive assessment of RNA-seq accuracy, reproducibility and information content by the Sequencing Quality Control Consortium. Nat Biotechnol 32, 903–914 (2014).

48. Eleveld, T. F., et al. In Silico Analysis of RAS-MAPK Activation in Neuroblastoma Identifies a Putative Feed Forward Loop Involving RET and SHC3, and a Possible Link between RAS-MAPK Signaling and EMT. https://dare.uva.nl (2019).

49. Verhoeven, B. M. et al. The immune cell atlas of human neuroblastoma. Cell Rep Med 3, (2022).

50. Calandra, T. & Roger, T. Macrophage migration inhibitory factor: a regulator of innate immunity. Nat Rev Immunol 3, 791–800 (2003).

51. Cavalli, E. et al. Emerging role of the macrophage migration inhibitory factor family of cytokines in neuroblastoma. Pathogenic effectors and novel therapeutic targets? Molecules vol. 25 Preprint at 10.3390/molecules25051194 (2020).

52. Guda, M. R., et al. Pleiotropic Role of Macrophage Migration Inhibitory Factor in Cancer. Am J Cancer Res vol. 9 www.ajcr.us/ (2019).

53. Noe, J. T. & Mitchell, R. A. MIF-Dependent Control of Tumor Immunity. Front Immunol 11, (2020).

54. Figueiredo, C. R. et al. Blockade of MIF-CD74 Signalling on Macrophages and Dendritic Cells Restores the Antitumour Immune Response Against Metastatic Melanoma. Front Immunol 9, (2018).

55. Mora Barthelmess, R., Stijlemans, B. & Van Ginderachter, J. A. Hallmarks of Cancer Affected by the MIF Cytokine Family. Cancers (Basel) 15, 395 (2023).

56. Thiele, M., Donnelly, S. C. & Mitchell, R. A. OxMIF: a druggable isoform of macrophage migration inhibitory factor in cancer and inflammatory diseases. J Immunother Cancer 10, (2022).

57. Cerezo-Wallis, D. et al. Midkine rewires the melanoma microenvironment toward a tolerogenic and immune-resistant state. Nat Med 26, 1865–1877 (2020).

58. Filippou, P. S., Karagiannis, G. S. & Constantinidou, A. Midkine (MDK) growth factor: a key player in cancer progression and a promising therapeutic target. Oncogene 2019 39:10 39, 2040–2054 (2019).

59. Tirosh, I. et al. Dissecting the multicellular ecosystem of metastatic melanoma by single-cell RNA-seq. Science (1979) 352, 189–196 (2016).

60. Bosse, K. R. et al. Identification of GPC2 as an Oncoprotein and Candidate Immunotherapeutic Target in High-Risk Neuroblastoma. Cancer Cell 32, (2017).

61. Pascual-Pasto, G. et al. GPC2 antibody-drug conjugate reprograms the neuroblastoma immune milieu to enhance macrophage-driven therapies. J Immunother Cancer 10, (2022).

62. Chirnomas, D., Hornberger, K. R. & Crews, C. M. Protein degraders enter the clinic - a new approach to cancer therapy. Nature Reviews Clinical Oncology vol. 20 265–278 Preprint at 10.1038/s41571-023-00736-3 (2023).

63. Dale, B. et al. Advancing targeted protein degradation for cancer therapy. Nat Rev Cancer 21, 638–654 (2021).

64. Xiao, Z. et al. Proteolysis Targeting Chimera (PROTAC) for Macrophage Migration Inhibitory Factor (MIF) Has Anti-Proliferative Activity in Lung Cancer Cells. Angewandte Chemie - International Edition 60, 17514–17521 (2021).

65. Kholosy, W. M. et al. Neuroblastoma and DIPG organoid coculture system for personalized assessment of novel anticancer immunotherapies. J Pers Med 11, (2021).

66. Bin, Q., Johnson, B. D., Schauer, D. W., Casper, J. T. & Orentas, R. J. Production of macrophage migration inhibitory factor by human and murine neuroblastoma. Tumor Biology 23, 123–129 (2002).

67. Lubetsky, J. B., Swope, M., Dealwis, C., Blake, P. & Lolis, E. Pro-1 of macrophage migration inhibitory factor functions as a catalytic base in the phenylpyruvate tautomerase activity. Biochemistry 38, 7346–7354 (1999).

68. Stosic-Grujicic, S., Stojanovic, I. & Nicoletti, F. MIF in autoimmunity and novel therapeutic approaches. Autoimmunity Reviews vol. 8 244–249 Preprint at 10.1016/j.autrev.2008.07.037 (2009).

69. Garcia-Gerique, L. et al. MIF/CXCR4 signaling axis contributes to survival, invasion, and drug resistance of metastatic neuroblastoma cells in the bone marrow microenvironment. BMC Cancer 22, (2022).

70. Cavalli, E. et al. Overexpression of Macrophage Migration Inhibitory Factor and Its Homologue D-Dopachrome Tautomerase as Negative Prognostic Factor in Neuroblastoma. Brain Sci 9, 284 (2019).

71. Cavalli, E. et al. Emerging Role of the Macrophage Migration Inhibitory Factor Family of Cytokines in Neuroblastoma. Pathogenic Effectors and Novel Therapeutic Targets? Molecules 25, 1194 (2020).

72. Ren, Y. et al. Upregulation of macrophage migration inhibitory factor contributes to induced N-Myc expression by the activation of ERK signaling pathway and increased expression of interleukin-8 and VEGF in neuroblastoma. Oncogene 23, 4146–4154 (2004).

73. Fetahu, I. S. et al. Single-cell transcriptomics and epigenomics unravel the role of monocytes in neuroblastoma bone marrow metastasis. Nat Commun 14, 3620 (2023).

74. Yaddanapudi, K. et al. MIF Is Necessary for Late-Stage Melanoma Patient MDSC Immune Suppression and Differentiation. Cancer Immunol Res 4, 101–112 (2016).

75. Tessaro, F. H. G. et al. Single-cell RNA-seq of a soft-tissue sarcoma model reveals the critical role of tumor-expressed MIF in shaping macrophage heterogeneity. Cell Rep 39, 110977 (2022).

76. Sullivan, P. M. et al. FGFR4-Targeted Chimeric Antigen Receptors Combined with Anti-Myeloid Polypharmacy Effectively Treat Orthotopic Rhabdomyosarcoma. Mol Cancer Ther 21, 1608–1621 (2022).

77. Li, W. et al. Identification of potential resistance mechanisms and therapeutic targets for the relapse of BCMA CAR-T therapy in relapsed/refractory multiple myeloma through single-cell sequencing. Exp Hematol Oncol 12, 44 (2023).

78. Sreeramkumar, V., Fresno, M. & Cuesta, N. Prostaglandin E _2_ and T cells: friends or foes? Immunol Cell Biol 90, 579–586 (2012).

79. Yan, X., Orentas, R. & Johnson, B. Tumor-derived macrophage migration inhibitory factor (MIF) inhibits T lymphocyte activation. Cytokine 33, 188–198 (2006).

80. Wang, X. et al. MIF Produced by Bone Marrow–Derived Macrophages Contributes to Teratoma Progression after Embryonic Stem Cell Transplantation. Cancer Res 72, 2867–2878 (2012).

81. Otvos, B. et al. Cancer Stem Cell-Secreted Macrophage Migration Inhibitory Factor Stimulates Myeloid Derived Suppressor Cell Function and Facilitates Glioblastoma Immune Evasion. Stem Cells 34, 2026–39 (2016).

82. Wang, D., et al. Significance of the vascular endothelial growth factor and the macrophage migration inhibitory factor in the progression of hepatocellular carcinoma. Oncol Rep 31, 1199–1204 (2014).

83. Kindt, N. et al. Macrophage migration inhibitory factor in head and neck squamous cell carcinoma: clinical and experimental studies. J Cancer Res Clin Oncol 139, 727– 737 (2013).

84. Xia, H. H. et al. Serum macrophage migration-inhibitory factor as a diagnostic and prognostic biomarker for gastric cancer. Cancer 115, 5441–5449 (2009).

85. Wu, L. et al. The level of macrophage migration inhibitory factor is negatively correlated with the efficacy of PD-1 blockade immunotherapy combined with chemotherapy as a neoadjuvant therapy for esophageal squamous cell carcinoma. Transl Oncol 37, 101775 (2023).

86. O’Reilly, C., Doroudian, M., Mawhinney, L. & Donnelly, S. C. Targeting MIF in Cancer: Therapeutic Strategies, Current Developments, and Future Opportunities. Med Res Rev 36, 440–460 (2016).

87. Kildisiute, G. et al. Tumor to normal single-cell mRNA comparisons reveal a pan-neuroblastoma cancer cell. Sci Adv 7, (2021).

88. Strijker, J. G. M. et al. αβ-T Cells Engineered to Express γδ-T Cell Receptors Can Kill Neuroblastoma Organoids Independent of MHC-I Expression. J Pers Med 11, 923 (2021).

89. Pscheid, R. et al. Three-Dimensional Modeling of Solid Tumors and Their Microenvironment to Evaluate T Cell Therapy Efficacy In Vitro. The Journal of Immunology 211, 229–240 (2023).

90. Xavier, D. et al. Heat ’n Beat: A Universal High-Throughput End-to-End Proteomics Sample Processing Platform in under an Hour. Anal Chem 96, 4093–4102 (2024).

