## Supplementary figures for "Blocking MIF secretion enhances CAR T-cell efficacy against neuroblastoma"

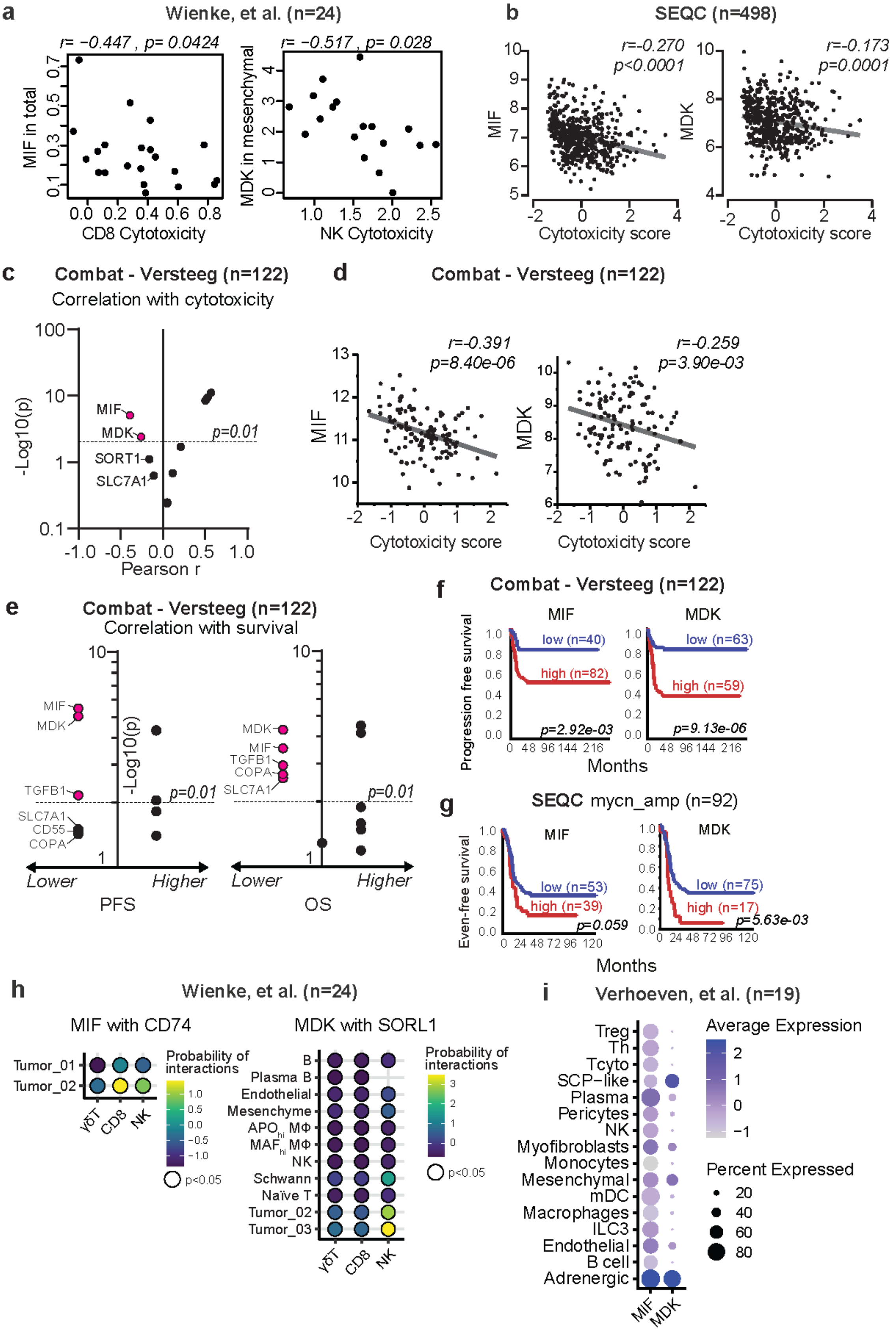

**Supplementary Figure 1: a,** Correlation of *MIF* and *MDK* with CD8 T cell cytotoxicity and NK Cytotoxicity in scRNAseq data[30]. **b,** Correlation of *MIF* and *MDK* with cytotoxicity score in bulk-RNAseq dataset of SEQC cohort consisting of 498 neuroblastoma tumors (r2.amc.nl/; Tumor Neuroblastoma - SEQC - 498 - RPM - seqcnb1; GSE49710)[47] **c,** Correlation of 13 selected genes from Fig. 1d with cytotoxicity in dataset with bulk-RNA data from 122 neuroblastoma tumors (r2.amc.nl/; Tumor Neuroblastoma (Combat) - Versteeg - 122 - MAS5.0(bc) - u133p2; GSE16476). **d,** Correlation of *MIF* and *MDK* with cytotoxicity score in bulk-RNAseq dataset of Combat-Versteeg cohort consisting of 122 neuroblastoma tumors. **e,** Survival analysis using the 13 selected genes from Fig. 1d using the Combat - Versteeg cohort. Left panel represents progression-free survival and right panel represents overall survival. **f**, Kaplan-Meier curve indicating event-free survival for high- or low expression of *MIF* (left panel, expression cutoff: 2026.1) and *MDK* (right panel, expression cutoff: 341.7) in Combat-Versteeg dataset. **g,** Kaplan-Meier curve indicating event-free survival for high- or low expression of *MIF* (left panel, expression cutoff: 206.358) and *MDK* (right panel, expression cutoff: 276.327) only for the MYCN-amplified subset from the SEQC dataset (n=92). **h,** Bubbelplot showing the probability of interaction between *MIF* on cell subsets on the y-axis and *MIF* receptor CD74 on immune cell subsets on the x-axis (left) and the probability of interaction between *MDK* on cell subsets on the y-axis and *MDK* receptor *SORL1* on immune cell subsets on the x-axis (right). Only significant values with a $p<0.05$ are shown. Data from the Wienke cohort has been used[30]. **i,** Validation of expression of *MIF* and *MDK* on cell subsets in single-cell RNAseq dataset of 19 patients published by Verhoeven, et al[49].

**a**

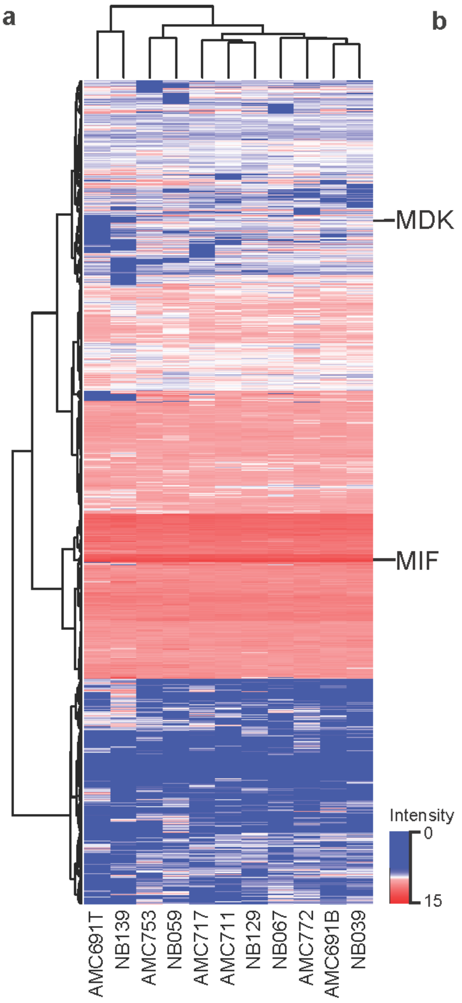


**b**

| Tumoroid | Treatment stage | *MYCN* status | *ATRX* status | Other |
|---|---|---|---|---|
| NB039 | Relapse | Amplified | Wild Type | 1p loss, 11q loss, 17q gain |
| AMC691B | Diagnosis | Amplified | Wild Type | 1p loss, 17q gain |
| AMC772 | Diagnosis | Non-Amplified | Homologous deletion | 11q loss, 17q gain |

**Supplementary Figure 2: a,** Heatmap showing the expression of a total of 5297 proteins in whole tumoroid lysates. MIF is the 20$^{th}$ most abundant protein with high expression in 10 out of 11 tumoroids and MDK as 3632$^{th}$ with expression in 8 out of 11 tumoroids. **b,** Table showing characteristics of three chosen tumoroids for secretome analysis.

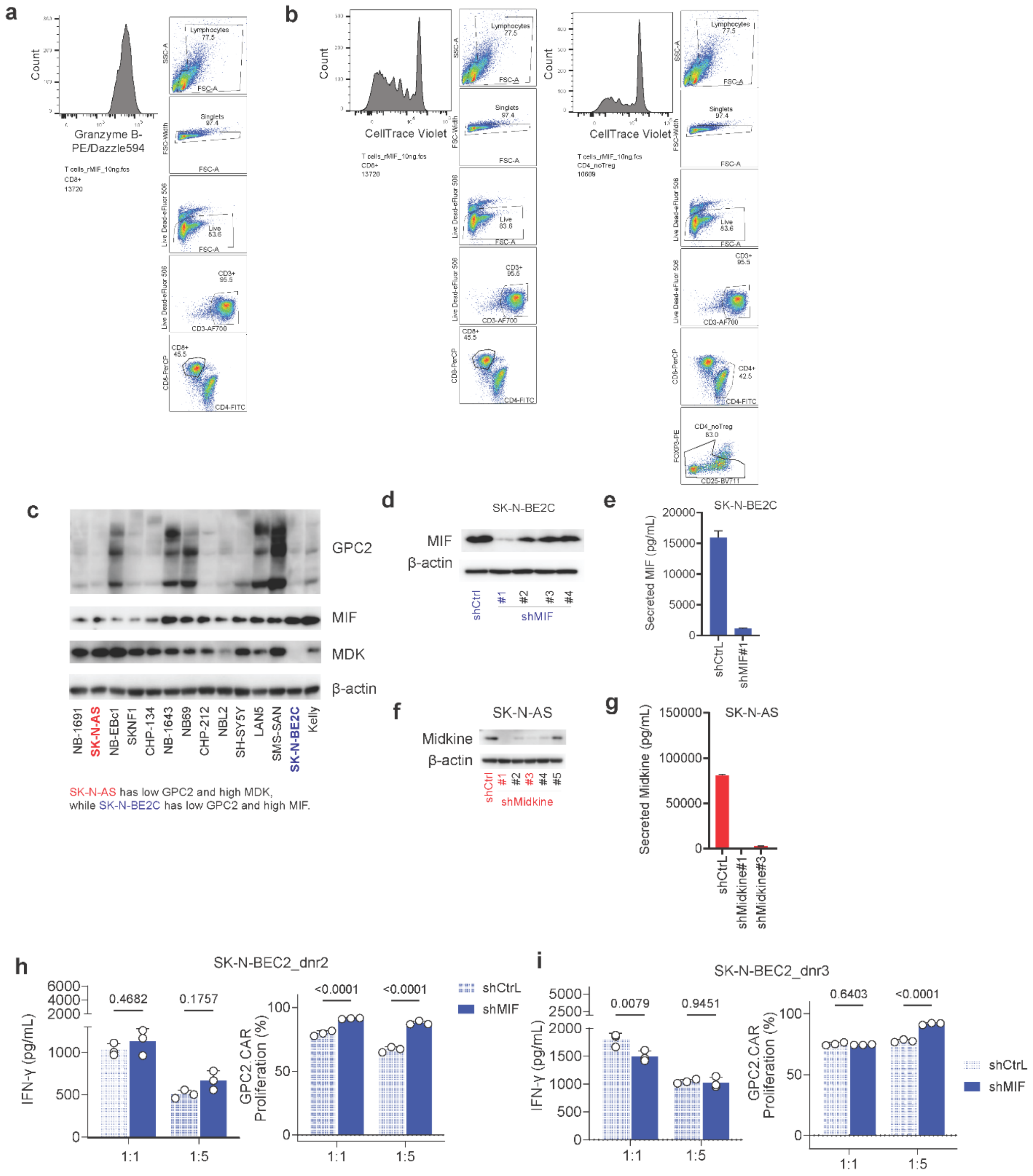


**Supplementary figure 3: a,** Gating strategy for Fig 3a. **b,** Gating strategy for Fig 3b. **c,** Western blot of 14 neuroblastoma cell lines for GPC2, MIF and MDK. One cell line with low GPC2 and high MIF and one line with low GPC2 and high MDK were chosen for shRNA modulation of MIF and MDK, respectively. β-actin was used as a loading control. **d,** Validation of shRNA knock-down by western blot for MIF on cell line with shCtrl or shMIF. shMIF construct #1 was chosen for further testing. β-actin is used as a loading control. **e,** Validation of shRNA knock-down of secreted MIF by MIF ELISA. **f,** Validation of shRNA knock-down by western blot for MDK on cell line with shCtrl or shMDK. shMDK constructs #1 and #3 were chosen for further testing. β-actin is used as a loading control. **g,** Validation of shRNA knock-down of secreted MDK by MDK ELISA. **h,i,** Activation of GPC2 CAR T-cells in co-culture with SK-N-BEC2 models as measured by IFN-γ ELISA (left panel) and GPC2 CAR T-cell proliferation (right panel)

in two effector : target ratios. shCtrl SK-N-BEC2 model (light blue) and shMIF SK-N-BEC2 (dark blue). Statistical analysis shows Two-way ANOVA with Šídák's multiple comparisons test. (*n=1* CAR donor with several technical replicates.

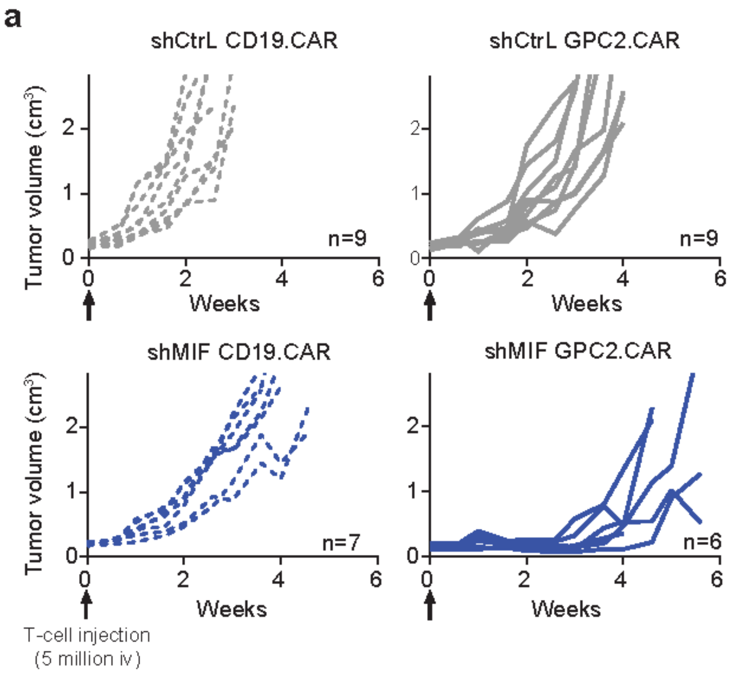


**Supplementary figure 4: a,** SK-N-BEC2 shCtrl or shMIF tumor growth. Measuring of tumor size started when CD19 or GPC2 CAR-T cells were injected ($5x10^6$ iv, at arrow indication). Experimental groups of n=6 – n=9.

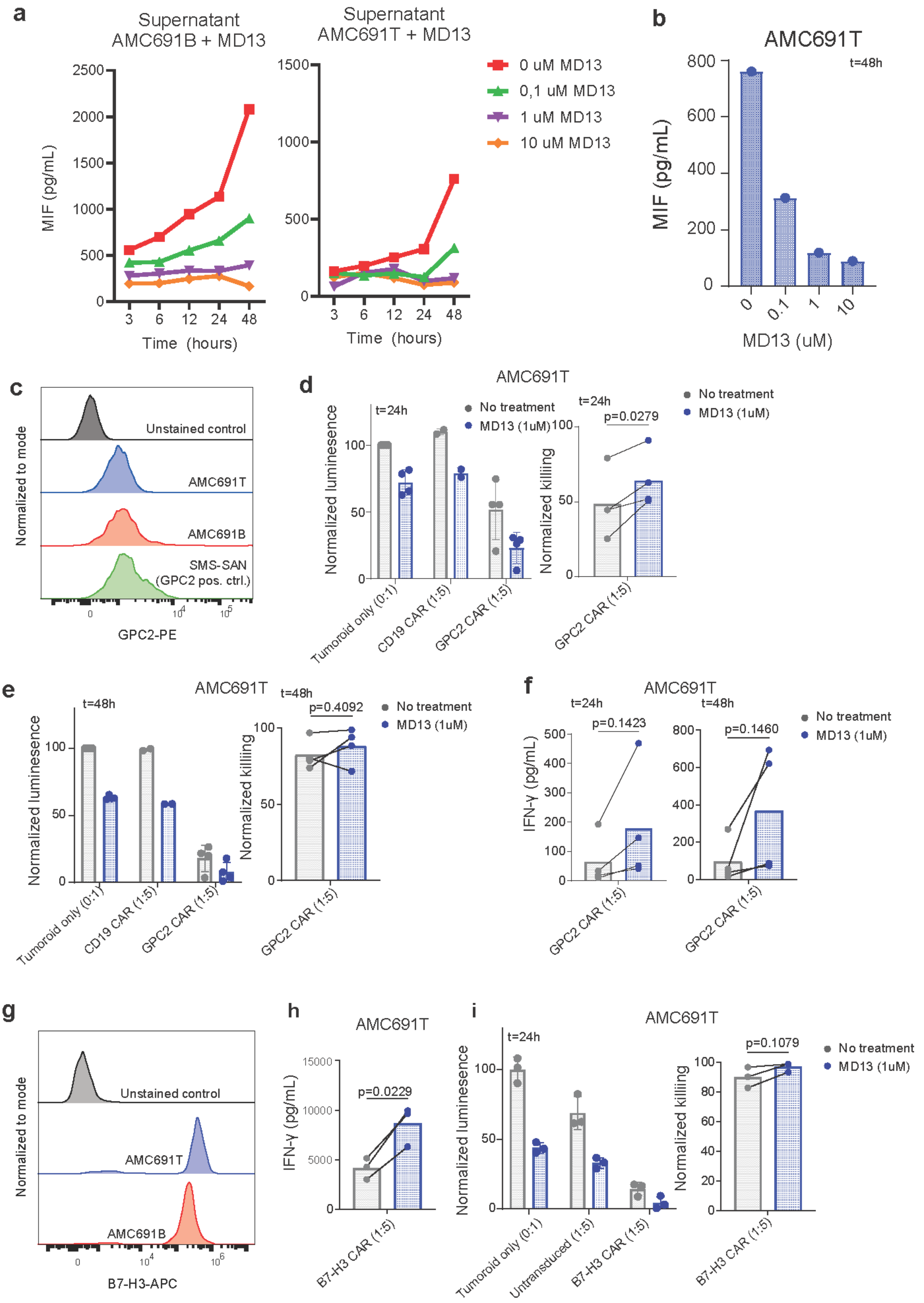


**Supplementary figure 5: a,** Concentration of MIF in culture supernatant after treatment with MIF PROTAC MD13. At t=0, medium was refreshed with medium containing MD13. MIF concentration was measured using Luminex. **b,** MIF concentration AMC691T tumoroid model 48 hours of treatment with

0.1, 1 or 10µM MD13. Measured using Luminex. **c,** Flow cytometry results of GPC2 staining on AMC691B (red) and AMC691T (blue). SMS-SAN (green) was used as a positive control with known GPC2 expression. **d,** Left panel: Luminescence signal of luciferase transduced tumoroid model AMC691T after co-culture of 24 hours. Normalized to untreated tumoroid only. Tumoroids were pre-treated for 48 hours before co-culture. Right panel: Normalized GPC2 CAR-T cell killing. Data were normalized to the tumoroid only untreated or treated control, respectively. Statistical analysis shows results for paired t-test. (*n=2* CD19-CAR T-cell donors, *n=4* GPC2-CAR T-cell donors). **e,** Left panel: Luminescence signal of luciferase transduced tumoroid model AMC691T after co-culture of 48 hours. Normalized to untreated tumoroid only. Tumoroids were pre-treated for 48 hours before co-culture. Right panel: Normalized GPC2 CAR-T cell killing. Data were normalized to the tumoroid only untreated or treated control, respectively. Statistical analysis shows results for paired t-test. (*n=2* CD19-CAR T-cell donors, *n=4* GPC2-CAR T-cell donors). **f,** IFN-γ concentration of supernatant from co-culture of AMC691T without treatment (grey) or with 1µM MD13 treatment (blue) in combination with GPC2 CAR-T cell in a 1:5 Effector : Target ratio, as measured by ELISA. Left panel shows concentration at 24 hours and right panel shows concentration at 48 hours. Statistical analysis shows results for paired t-test. (*n=4* GPC2-CAR T-cell donors). **g,** Flow cytometry analysis of B7-H3 staining on AMC691B and AMC691T. Unstained control is AMC691T. **h,** IFN-γ concentration of supernatant from co-culture of AMC691T without treatment (grey) or with 1µM MD13 treatment (blue) in combination with B7-H3 CAR-T cell in a 1:5 Effector : Target ratio, as measured by ELISA. (*n*=3 B7-H3-CAR T-cell donors). **i,** Left panel: Luminescence signal of luciferase transduced tumoroid models after co-culture of 24 hours. Normalized to untreated tumoroid only. Tumoroids were pre-treated for 48 hours before co-culture. Right panel: Normalized B7-H3 CAR-T cell killing. Data were normalized to the tumoroid only untreated or treated control, respectively. Statistical analysis shows results for paired t-test. (*n=3* CAR T-cell donors with untransduced controls).
